# Systematic screening identifies elevated neurovascular BCL-XL in Parkinson’s disease

**DOI:** 10.64898/2026.05.03.722488

**Authors:** Kreesan Reddy, Eden Yin, Xi Hua Wu, Taylor Stevenson, Brigid Ryan, Helen C. Murray, Deborah Young, Richard L. M. Faull, Maurice A. Curtis, Glenda M. Halliday, Ronald Melki, Mike Dragunow, Birger Victor Dieriks

## Abstract

**Background:** Vascular cells are emerging as active players in Parkinson’s disease (PD), yet their molecular contribution to α-Synuclein (α-Syn) pathology remains undefined. Here, we show that human brain pericytes respond in a distinct manner to unique α-Syn strains, with systematic validation identifying BCL-XL as a potential regulator of the neurovascular unit in PD.

**Methods:** Primary human brain-derived pericytes were exposed to five recombinant α-Syn strains (Fibrils, Fibrils-65, Fibrils-91, Fibrils-110, and Ribbons). Transcriptomic profiling identified differentially expressed genes (DEGs), which were validated at the protein level using multiplex immunocytochemistry and *in situ* labelling of post-mortem middle temporal gyrus (MTG) tissue microarrays from PD (n = 24) and neurologically normal (n = 24) cases.

**Results:** α-Syn strain exposure produced 300 DEGs with limited overlap between strains. BCL-XL and CSNK1D were upregulated in α-Syn-treated pericytes. In post-mortem PD tissue BCL-XL showed marked pericyte-specific elevation in the MTG and increased pericytic and microglial expression in the substantia nigra.

**Conclusion:** BCL-XL emerges as a potential regulator of pericyte and microglial resilience in PD, linking acute α-Syn strain–specific responses in pericytes to broader neurovascular alterations. Its upregulation likely represents a generalised compensatory response to chronic α-Syn–associated stress beyond individual strain effects, identifying BCL-XL as a possible therapeutic target within the neurovascular unit.

## Background

Pericytes are increasingly recognised as active contributors to Parkinson’s disease (PD) pathology, extending beyond their classical role in maintaining blood–brain barrier integrity [1]. These contractile vascular cells line capillary walls and are essential for vascular stability, immune regulation, and neurovascular coupling [2]. Evidence indicates that pericyte dysfunction occurs early in PD and can amplify neuroinflammation, compromise barrier integrity, and heighten neuronal vulnerability. In post-mortem PD brains, pericytes show altered morphology and inflammatory phenotypes, while experimental models demonstrate that pericyte loss or activation exacerbates neurodegeneration [3,4]. Despite growing recognition of vascular involvement, the molecular mechanisms driving pericyte responses in PD remain poorly defined.

α-Synuclein (α-Syn) aggregation is the defining molecular hallmark of PD, driving progressive neurodegeneration through the formation and spread of pathological inclusions [5–8]. α-Syn assembles into structurally distinct fibrillar strains with unique biochemical and pathological properties. These conformational variants differ in seeding efficiency, cell-type tropism, and regional distribution, contributing to the clinical and pathological heterogeneity of synucleinopathies [9–20]. Importantly, α-synucleinopathic disease phenotypes are shaped not only by intrinsic α-Syn strain characteristics but also by cell type–specific factors, regional localisation, and the broader macroenvironment [21]. Given the presence of α-Syn pathology within the neurovascular unit and the capacity of pericytes to internalise and degrade α-Syn aggregates [22–24], understanding how pericytes respond to strain-specific α-Syn species is critical for defining neurovascular mechanisms in PD progression.

In this study, primary human brain-derived pericytes were exposed to pure recombinant α-Syn strains, followed by integrated transcriptomic and immunocytochemical analyses, with extensive protein-level validation in post-mortem PD tissue. Transcriptomic profiling revealed that exposure to distinct α-Syn strains induces distinct gene expression programs in pericytes, providing a foundation for subsequent *in situ* validation and functional interrogation. Notably, while strain-specific responses were evident *in vitro*, tissue-level analyses suggested that cellular context and the diseased macroenvironment may supersede or modulate intrinsic strain differences. Among the identified targets, the anti-apoptotic protein BCL-XL was consistently upregulated both *in vitro* and *in situ*, localising to pericytes and microglia in PD brain tissue. These findings implicate BCL-XL as a potential regulator of neurovascular resilience in PD, linking acute strain-specific responses in pericytes to a broader compensatory response to chronic α-Syn–associated stress within the neurovascular unit.

## Methods

### Study Design

To identify α-Syn strain-specific targets in PD, a sequential multi-stage validation approach was undertaken. Primary human brain-derived pericytes, herein referred to as *pericytes*, were isolated from individuals with PD (n = 3) and neurologically normal controls (n = 3). These mural cells, located along the abluminal surface of capillaries and involved in blood–brain barrier regulation and vascular stability, were treated with five distinct α-Syn aggregate strains (Fibrils, Fibrils-65, Fibrils-91, Fibrils-110, and Ribbons [9,10,12]) or either a monomeric α-Syn control or a phosphate buffered saline (PBS, pH 7.4) no-treatment control for four hours, followed by a media change and a subsequent 20-hour incubation. The brief exposure period allowed aggregate binding to the cell membrane and subsequent endocytosis, while minimising excessive intracellular accumulation [24]. RNA sequencing was then performed to identify differentially expressed genes (DEGs). A subset of DEGs was selected for protein-level validation based on fold-change, significance, and commercial antibody availability. Protein expression was assessed in pericytes using the same treatment paradigm as in the transcriptomic experiments, enabling direct comparison between RNA and protein expression levels through immunocytochemical labelling of treated and control cells.

In parallel, the selected antibodies were evaluated in formalin-fixed, paraffin-embedded human MTG tissue. Labelling protocols were optimised iteratively by varying antigen retrieval (citric or Tris EDTA buffer with or without formic acid) and antibody concentrations, and evaluated for signal intensity, specificity, and lipofuscin interference. Antibodies showing consistent, high-quality labelling were retained to map cell type-specific expression in human tissue, with negative controls included to evaluate background fluorescence and non-specific binding.

Antibodies demonstrating robust immunoreactivity in MTG tissue were subsequently applied to tissue microarrays (TMAs) containing samples from 24 PD and 24 neurologically normal cases, allowing efficient high-throughput screening of candidate DEGs at the protein level. Each TMA was multiplex-labelled with up to six antibodies and imaged using a Zeiss high-throughput slide scanner. Proteins showing differential expression profiles were then examined across additional brain regions (MTG, substantia nigra, medulla, middle frontal gyrus, and cerebellum) from 10 PD and 10 neurologically normal cases representing progressive stages of disease, with supplementary western blotting performed to confirm antibody specificity [8,25].

Given that DEGs were identified following α-Syn exposure, confocal microscopy was used to determine whether candidate protein changes co-localised with α-Syn pathology. Because the phosphorylated serine 129 (pS129) antibody alone does not detect all inclusions, two complementary antibodies were used: #849101, targeting residues 34–35 of the N-terminus, and #ab184674, recognising the pS129 epitope [26,27]. This stepwise workflow refined the transcriptomic dataset to a focused group of proteins validated *in vitro* and in human tissue, confirming their altered expression at the protein level.

### *α*-Syn strain generation

Human wild-type monomeric α-Syn was expressed in E. coli BL21 DE3 CodonPlus cells (Agilent Technologies) and purified as previously detailed [28]. Subsequent generation of distinct fibrillar aggregates was then performed as done previously [9,10,12,24,29]. Quantification of the endotoxin levels of all preparations was carried out as detailed before [10,30], using a Pierce LAL Chromogenic Endotoxin Quantification kit to ensure all levels were below 0.02 endotoxin units/mg.

### Human Tissue

Post-mortem human brain tissue (Supplementary Table 1) was obtained from the Neurological Foundation Human Brain Bank at the University of Auckland with informed consent from donors or their next of kin. All samples underwent neuropathological assessment to confirm diagnosis. Neurologically normal control cases showed no neurological abnormalities, while PD cases exhibited clinical features of Parkinsonism and α-Syn pathology in the substantia nigra and frontal lobe, accompanied by characteristic neuronal and pigment loss. Ethical approval was granted by the University of Auckland Human Participants Ethics Committee (2014/011654).

### Human brain-derived pericyte isolation and culturing

Pericyte cultures were sourced from the Hugh Green Biobank (Supplementary Table 1), specifically from the middle temporal gyrus of post-mortem brains from individuals with PD or neurologically normal controls. The time between death and tissue collection had little impact on pericyte viability compared with factors such as age and disease state, with viable cultures obtained up to 48 hours post-mortem [24,31]. Detailed protocols for isolating and culturing human brain-derived pericytes relevant to this study have been previously described [31].

Tissue from the middle temporal gyrus was mechanically dissected and dissociated, followed by enzymatic digestion in Hank’s balanced salt solution (HBSS; Ca2+ and Mg2+ free; #14175095, ThermoFisher Scientific) containing 2.5 U/mL papain (#LS003124, Worthington Biochemical) and 100 U/mL DNase 1 (#18047019, ThermoFisher Scientific) for 30 minutes at 37°C with gentle rotation, including gentle titration after 15 minutes. The enzymatic digestion was halted by adding complete media (DMEM: F12; #11320033, ThermoFisher Scientific). Cells were collected by centrifugation (170g for 10 minutes), re-suspended in complete media, and plated in uncoated T75 flasks (75 cm^2^; EasYFlask™ with Nunclon™ Delta surface; #156499, Nunc). The cultures were incubated at 37°C with 5% CO2 until ready for experiments, grown in DMEM:F12 supplemented with 10% foetal bovine serum (FBS; #MORFBSFAU, Moregate) and 1% Penicillin/Streptomycin (#15140122, ThermoFisher Scientific). Pericytes used in this study were at passages 5– 9, free from astrocyte or microglial contamination [32,33]. Immunocytochemical analysis of late passages (beyond P4) confirmed expression of pericyte markers such as PDGFRβ, neural/glial antigen 2 (NG2), CD13, CD146, and desmin, consistent with human brain capillary-associated pericyte profiles[34].

### Pericyte *α*-Synuclein strain treatment

Cells were harvested for experiments by adding 2 mL 0.25% Trypsin-1mM ethylenediaminetetraacetic acid (EDTA; #25200056, ThermoFisher Scientific) and incubated for 2-5 minutes at 37°C to allow for cell detachment. Cells were collected in warm DMEM:F12 media. 10 μL of 1:1 Trypan Blue (#15250061, ThermoFisher Scientific) and cell suspension was prepared and added to a hemocytometer to allow for cell counting. Cells were re-suspended in the correct volume of DMEM:F12 to achieve the required cell density of 5,000 cells/well in a 96-well plate (#243656, Nunc). Pericytes (nCtrl = 3, nPD = 3) were treated at 37°C with the various pure α-Syn strains (Fibrils, Fibrils-65, Fibrils-91, Fibrils-110, Ribbons, 100 nM in PBS) or either a monomeric α-Syn control or a PBS (no treatment control) for 4 hours, washed and subsequently incubated for 20 hours. Cells were fixed in 4% paraformaldehyde.

## RNAseq

### Extraction and quality assessment of RNA

Total RNA was extracted using the RNeasy Minikit (Qiagen, #71104) following standard protocols. RNA quality and concentration were evaluated with the Agilent 2100 Bioanalyser using an RNA nanochip (Agilent Technologies) as per the manufacturer’s guidelines, confirming sufficient RNA yields in our samples. DV_200_ values were determined from the same Bioanalyser traces by applying the smear analysis function within the Agilent 2100 Expert Software (Agilent Technologies). RNA-seq libraries were prepared with the Truseq RNA Access kit, generating libraries selectively depleted of ribosomal RNA, as well as intronic and intergenic regions. These libraries underwent cluster generation and were sequenced on the Hiseq 2500 Ultra-High Throughput Sequencing System (Illumina) at the New Zealand Genomics Limited (NZGL) sequencing facility. The RNA-seq data can be viewed online in the Gene Expression Omnibus (GEO) database with the accession number (*to be uploaded*).

### Differential gene expression analysis

RNA sequencing of pericytes treated with α-Syn strains was performed, yielding paired-end reads in FASTQ format. The raw sequencing data were processed to remove adapter sequences, low-quality reads, and polyA tails. Data analysis followed the pipeline established by Trapnell et al. [35,36], employing Tophat2 (Version 2.1.0) and Bowtie2 (Version 2.2.6) for sequence alignment against the reference genome. Pre-processing of raw paired-end reads was conducted using Trimmomatic (Version 0.33) to eliminate overrepresented Illumina adapters, polyA sequences, and low-quality bases. The filtered reads were then aligned to the human reference genome (UCSC build hg19) using Tophat2 and Bowtie2. Assembled reads were subsequently processed with the Cufflinks suite (Version 2.2.1), which includes Cufflinks, Cuffmerge, and Cuffdiff, to identify DEGs between α-Syn strain-treated and PBS-treated pericytes. The assembled transcript data and mapped reads were integrated with the reference genome before being analysed for differential gene expression using Cuffdiff, producing quantified results for genes presented in Fragments per Kilobase of transcript per Million mapped reads (FPKM) values. DEGs were filtered by removing genes with a mean fold change value of less than 2 and greater than 0.5. Additional filtering to remove genes skewed by low expression values was performed by firstly removing any genes in which 2 or more pericyte cases had a 0 FPKM value for both treated and control groups, and secondly by removing genes in which 3 or more cases had FPKM values less than 1. A final manual check was performed to remove any genes that had more than one case overlapping the FPKM values of the other treatment group. The final list of genes was considered statistically significant (p < 0.05) and differentially expressed.

### Immunocytochemical Staining and Imaging

Immunocytochemical staining of fixed pericytes was done as previously described, using primary and secondary antibodies listed in Supplementary Tables 2-4 [22,23]. Imaging was carried out using an ImageXpress® Micro XLS automated fluorescence microscope (Molecular Devices) housed within the Biomedical Imaging Research Unit at the University of Auckland. The system was equipped with a 20× (0.45 NA) CFI Super Plan Fluor ELWD ADM objective lens and a Lumencor Spectra X light engine. Autofocus settings and exposure times were optimised for each plate to account for variations in antibody performance and plate manufacturing, ensuring consistent image quality.

Initial validations of DEGs at the protein level were carried out in neurologically normal (n = 3) and PD (n = 3) pericytes. To assess protein expression, cells were treated with the α-Syn strain that had induced the corresponding DEG.

### Immunohistochemical Staining and Imaging

Immunohistochemical staining was carried out as previously described, using 7 µm-thick formalin-fixed paraffin-embedded tissue microarrays and whole tissue sections [22,37–39]. A detailed description of the staining protocol is available at:https://dx.doi.org/10.17504/protocols.io.5qpvo3wdzv4o/v1

Tissue information is listed in Supplementary Table 1. Primary antibodies used to label tissue are detailed in Supplementary Table 2-3, with the antigen retrieval methods optimised to each antibody (Supplementary Table 5). Secondary antibodies used are listed in Supplementary Table 4. To ensure the specificity of BCL-XL labelling, additional validation of the BCL-XL antibody was performed as detailed in Supplementary Methods [40].

Images of stained tissue sections were acquired using a Zeiss Axio Imager.Z2 microscope with a Colibri 7 illumination system. Hardware control, slide scanning, and image stitching were performed using Metafer 5 (version 4.4.114) and VSlide (version 2.0.114) software at the University of Sydney node of Microscopy Australia. Scans were captured with FLUAR 5×/0.25 and Plan Apochromat 20×/0.8 objectives, with exposure settings for each fluorescent channel optimised individually for every section.

Confocal imaging was performed using an Olympus FV4000 confocal microscope (Evident) with a 60x (1.42 NA) UPlanxApo oil immersion objective lens, Next-Generation SilVIR detectors coupled with a Marzhauser motorised stage. Images were captured using the Olympus CellSens FV imaging software and exported as raw .oir files before processed in Fiji (version 1.54k).

## Image Analysis

### Immunocytochemistry Image and Statistical Analysis

Image analysis was performed using MetaXpress™ v5.3.04 (Molecular Devices). For each experimental paradigm, a unique module was created to enable quantification of specific protein labelling using the required analysis metric. Mean and integrated intensity was used as a measure of protein expression. To quantify the mean and integrated intensity, background labelling was removed by setting a threshold value that served as the lower limit for positive staining values. After background fluorescence was removed, a size filter was applied to remove debris and non-specific staining. Subsequent detection of positive staining was then measured and normalised to the number of Hoechst-labelled nuclei to account for cell number effects on protein expression. Data were exported in .csv format for subsequent statistical analysis using R (Version 4.5.1). To compare expression differences between strain treated and PBS control pericytes datasets were first tested for normality using the Shapiro-Wilk test. Subsequent analysis using a Paired t-test or Wilcoxon Signed-Rank test was conducted for parametric and non-parametric data, respectively. Results were considered significant if p < 0.05. In addition, Cohen’s d was calculated to assess the magnitude of observed effects independent of sample size, enabling standardisation of effect sizes to determine whether changes in protein expression across pericyte lines were consistent. Cohen’s d values of 0.2, 0.5, and 0.8 were considered small, medium, and large effects, respectively.

Figures were generated using a combination of R, Fiji (Version 1.54k), and the Adobe Creative Suite (Version 6.7.0.278).

### Immunohistochemistry Image and Statistical Analysis

Immunohistochemical image analysis was performed in a multi-step manner in Fiji (Version 1.54k). Exported images were manually reviewed to exclude those with blurred or damaged tissue. The remaining images underwent a three-step workflow to quantify tissue area, background signal intensity, and specific labelling load. Tissue areas were outlined manually using the selection and measure tool in Fiji (Version 1.54k). For the quantification of background signal intensity for each marker, specific protein staining was segmented using the rolling ball radius method, which was optimised for each image series. After removing background noise, the Triangle auto-thresholding function was applied to finalise the segmentation of specific staining. The specific staining was then subtracted from the original image to measure the residual background tissue signal. Protein load was quantified similarly by segmenting specific staining, as described earlier. This method was also used for channels containing known cell-type markers for neurons, astrocytes, pericytes, microglia and endothelial cells). Segmented images were further processed using the Image Calculator’s “And create” function, generating images showing co-localisation between the protein of interest and the cell marker. These segmented images were used to create a selection mask, which was applied to the original protein image to quantify the co-localised staining area. Load values were calculated by normalising the co-localised staining area to the total cell marker-stained area. To ensure cell-specific quantification was unbiased by differential expression in total cell marker staining, total cell marker area was normalised to total tissue area, confirming no significant differences in normalised cell marker between neurologically normal and PD tissues. Quantification of the whole section area staining was normalised to the total tissue area.

The Shapiro-Wilk test was used to assess the normality of data for each antibody stain. Depending on the distribution, either a t-test or Mann-Whitney U test was used to compare the healthy control and PD groups. Spearman correlations were used to assess relationships between variables, with statistical significance set at p < 0.05. Multiple comparisons correction was applied using the Benjamini-Hochberg method with a false discovery rate (FDR) set at 0.05. Correlation strength was categorised as follows: very weak (0.00–0.19), weak (0.20–0.39), moderate (0.40–0.59), strong (0.60–0.79), and very strong (0.80–1.00). Only correlations with a Spearman rho value above 0.59 and an adjusted p-value < 0.05 were retained for interpretation. To maintain relative differences when plotting data from multiple proteins or regions within the same graph, data were scaled using the scale function in R (Version 5.4.1). Graphs and representative images were created using the Adobe Creative Suite (Version 6.7.0.278) and the R ggplot2 package (Version 3.4.4).

## Results

### Differentially expressed genes vary based on strain treatment

RNAseq analysis of α-Syn strain-treated pericytes revealed 300 unique putative DEGs when assessing gene expression fold changes between control and treated pericytes, with minimal overlap occurring between treatment groups (Figure 1A). A subset of 256 DEGs were associated with only one strain treatment; Fibrils: 76 DEGs, Fibrils-65: 51 DEGs, Fibrils-91: 53 DEGs, Fibrils-110: 43 DEGs, Ribbons: 33 DEGs (Figure 1B). The remaining 44 were associated with at least two treatments, with no overlap occurring between all treatments (Figure 1C). DEGs for further validation were selected based on fold change, the significance of the change and the overlap of DEGs with the Developmental Studies Hybridoma Bank (DSHB) protein antibody database (Figure 1D). This filtering identified 37 unique DEGs as candidates for protein-level validation (Figure 1D, Supplementary Table 2).

**Figure 1.**
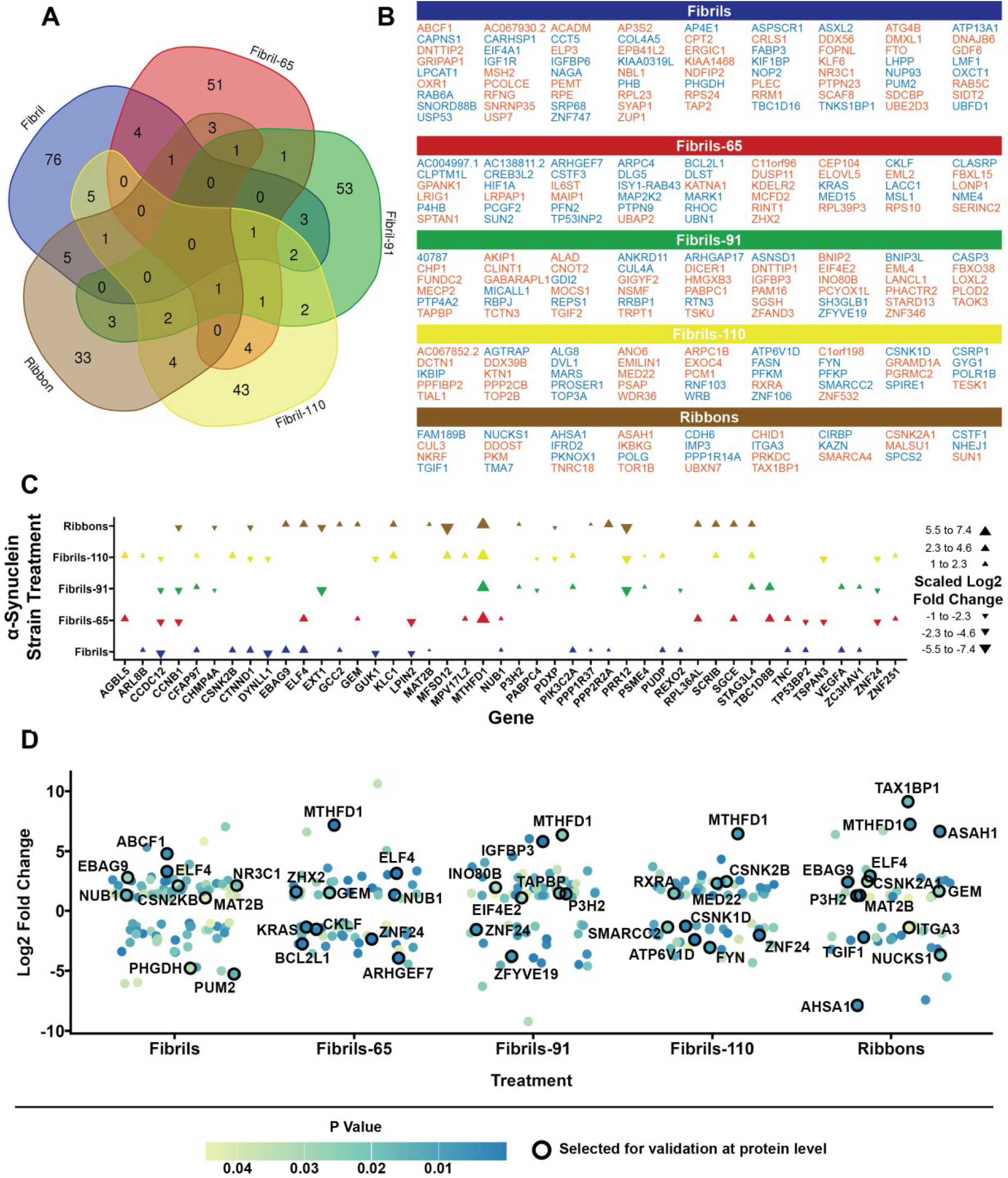
Differentially expressed genes vary based on unique *α*-Syn strain treatment. **(A)** Minimal overlap of differentially expressed genes across α-Syn strain treatments. **(B)**. Differentially expressed genes associated with only one α-Syn strain treatment. Orange text indicates upregulated genes, while blue text represents downregulated genes. **(C)**. Differentially expressed genes linked to two or more strain treatments. **(D)** Log2 Fold change of gene expression for each α-Syn strain treatment. Named points represent differentially expressed genes selected for validation based on Log2 Fold change, P value and antibody availability. Individual points were jittered to improve gene name readability.

### Protein validation in *α*-Syn strain-treated pericytes

Of the 43 targets assessed, six proteins showed reliable immunolabelling with signal intensities significantly above background autofluorescence (Figure 2A–F; Supplementary Figure 2A–F). Nuclear localisation was observed for BCL-XL (Encoded by *BCL2L1*), CSNK1D, EBAG9, and NUCKS1, while ITGA3 and MTHFD1 exhibited predominantly cytoplasmic staining. Quantitative analysis revealed increased expression of BCL-XL (1.04-fold; n = 6, p = 0.0075) and CSNK1D (1.25-fold; n = 6, p = 0.0294) in treated pericytes (Figure 2G).

**Figure 2.**
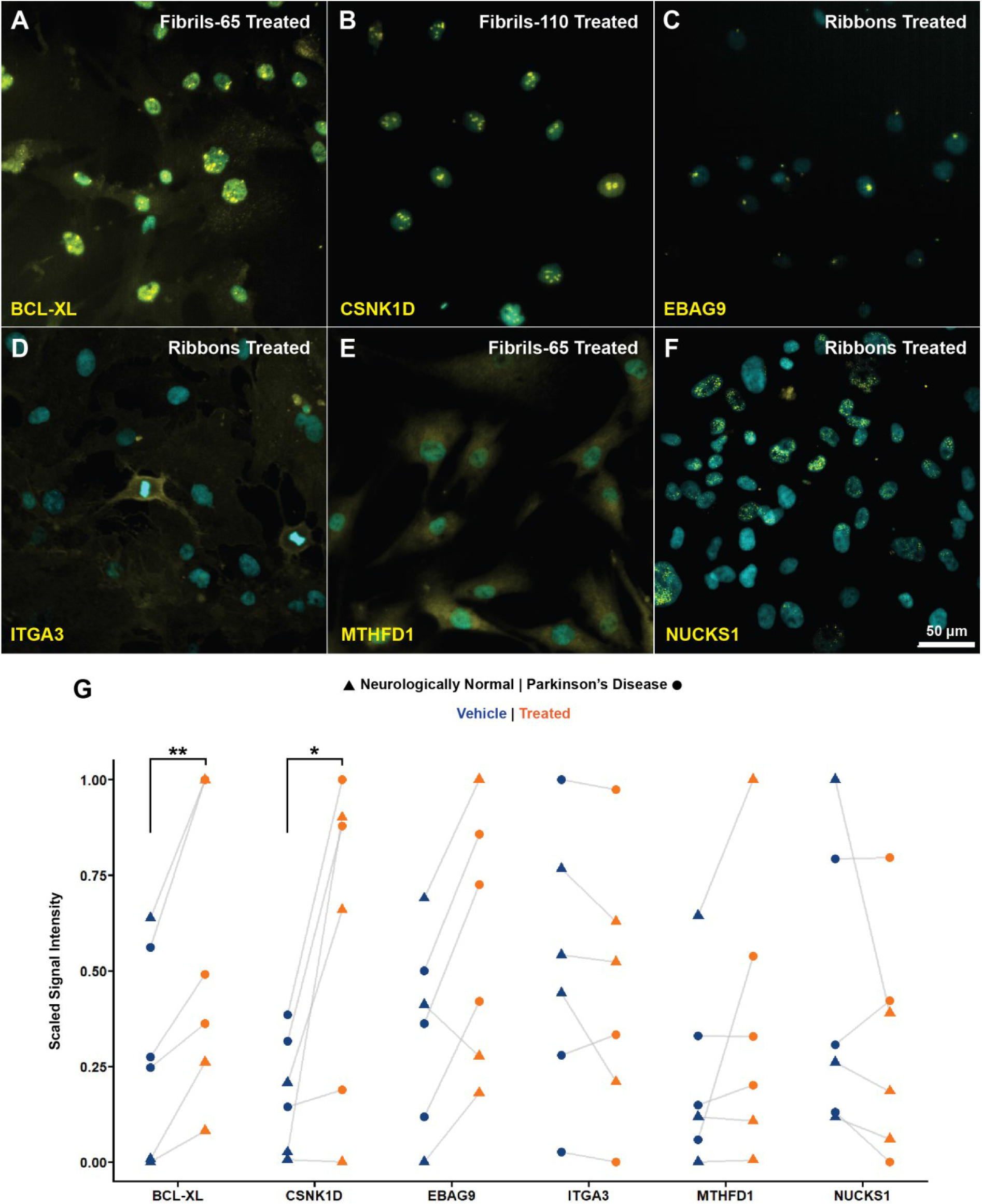
Immunocytochemical validation of differentially expressed gene analogous protein expression. **(A-F)** Representative images of differentially expressed genes’ analogous protein expression in pericytes. Scale bar represents 50 µm. **(G)** Quantification of protein expression in α-Syn strain-treated pericytes derived from neurologically normal (nCtrl = 3) and PD (nPD = 3) brains. BCL-XL expression increased by 1.04-fold (p = 0.0075), and CSNK1D by 1.25-fold (p = 0.0294) following α-Syn strain treatment. p < 0.05 (*), p < 0.01 (**)

Although the absolute fold changes were modest, effect size analysis demonstrated large and consistent treatment effects across cases (Cohen’s d: BCL-XL = 1.77; CSNK1D = 1.23), reflecting low inter-case variability and indicating that the observed increases were reproducible rather than driven by biological noise. Independent replication of the experimental paradigm confirmed this effect (Supplementary Figure 2G).

Comparison with RNA-seq findings revealed partial concordance between transcriptomic and protein-level responses. *ITGA3* and *EBAG9* demonstrated directional alignment, with ITGA3 showing reduced protein abundance (0.94-fold) consistent with its negative RNA-seq log_2_ fold change (−1.40 fold), and EBAG9 exhibiting increased protein expression (1.12-fold) in line with positive transcriptional regulation (+2.42-fold). In contrast, several targets displayed transcript–protein divergence. *BCL2L1* transcripts were markedly downregulated at the RNA level (−2.76 fold), despite a reproducible increase in BCL-XL protein expression. Similarly, *CSNK1D* showed transcriptional downregulation (−1.28 fold) while protein abundance increased. *MTHFD1* demonstrated strong transcriptional upregulation (+7.18-fold) but only a modest protein increase (1.08-fold), and *NUCKS1* transcripts were reduced (−3.68 fold) without a corresponding change in protein levels (0.99-fold).

### *In situ* validation of strain-specific proteins

#### Antibody optimisation

Analogous protein expression of DEGs was also assessed in post-mortem brain tissue from neurologically normal and PD cases to determine whether *in vitro* findings were recapitulated within a complex, chronic human disease environment encompassing heterogeneous α-Syn conformations. Prior to quantitative analysis, all antibodies were validated using MTG tissue from the region from which pericyte RNAseq samples were derived to determine their suitability for further investigation. All 37 antibodies were tested across a range of concentrations and antigen retrieval methods (Supplementary Table 5). Antibodies that were unsuccessful after several rounds were removed in addition to those producing a labelling signal, which was confounded by autofluorescence. This process resulted in 10 antibodies being selected for total and cell type-specific quantification using MTG TMAs, with multiplex panels tailored to the observed labelling patterns.

#### Tissue Microarray Screening

Of the 10 proteins analysed (ABCF1, ASAH1, BCL-XL, CSNK2B, FYN, GEM, MTHFD1, NUCKS1, PUM2, SMARCC2), cell-specific markers confirmed pericytic co-localisation for seven proteins (ASAH1, BCL-XL, FYN, GEM, MTHFD1, NUCKS1, SMARCC2, (Figure 3A-H, Supplementary Figure 2-3). Quantification of total and cell-specific protein labelling showed differential expression of BCL-XL and NUCKS1 between PD and neurologically normal cases (Figure 3I-M). Pericytic BCL-XL expression was elevated in PD cases, with a 2.58-fold increase compared to neurologically normal cases (*31*.*2 (nPD = 18) vs. 12*.*1 (nCtrl = 17); p = 0*.*0001*). Nuclear BCL-XL expression was also increased in PD (*1*.*75-fold, 7*.*7 vs. 4*.*4; p = 0*.*034*). In PD, neuronal NUCKS1 expression was reduced by 1.15-fold (*70*.*5, (nPD = 14) vs 81*.*3, (nCtrl) = 13; p = 0*.*006*). No differential expression was observed in total, astrocytic (ASAH1, BCL-XL, CSNK2B, GEM, MTHFD1, NUCKS1, PUM2), endothelial (ABCF1, ASAH1, FYN, GEM, MTHFD1, NUCKS1, PUM2, SMARCC2) or microglial (ASAH1, BCL-XL, FYN) co-localisation for all proteins tested (Supplementary Figure 4, Supplementary Table 7).

**Figure 3.**
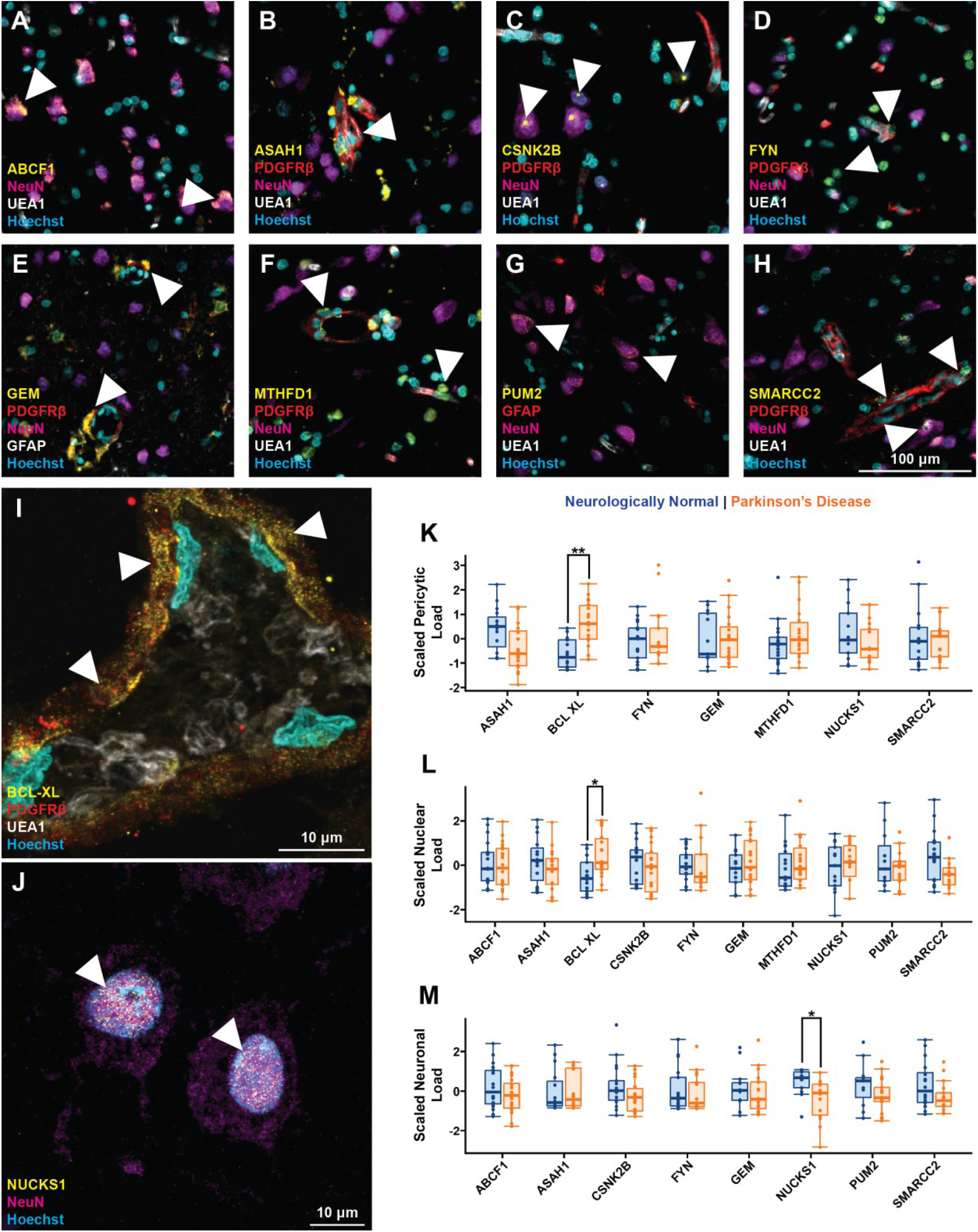
*In situ* expression of differentially expressed genes and analogous proteins. **(A-H)**. Expression of DEG analogous proteins in postmortem PD human MTG tissue. White arrows indicate points of specific staining. **(I)** Confocal Z-stack maximum projection showing BCL-XL (White arrows) co-localised with the pericyte marker PDGFRβ in PD MTG tissue. **(J**) Confocal Z-stack maximum projection showing NUCKS1 (white arrows) co-localised with the neuronal marker NeuN in PD MTG tissue. **(K)** Pericytic BCL-XL levels were higher in PD (2.58-fold; nPD = 18, nCtrl = 17, p = 0.0001). **(F)** Nuclear BCL-XL also increased in PD (1.75-fold, p = 0.034). (G) Neuronal NUCKS1 levels were lower in PD (1.15-fold; nPD = 14, nCtrl = 13, p = 0.006). Statistical analysis was performed using the Shapiro –Wilk test to assess normality, followed by t-tests or Mann–Whitney U tests as appropriate. P-values were corrected for multiple comparisons using the Benjamini–Hochberg method (FDR = 0.05). Load (protein area normalised to tissue area) datasets were scaled using thescale function in R to preserve relative differences when creating boxplots. Significance levels: p < 0.05 (*) and p < 0.01 (**).

Notably, SMARCC2 was not differentially expressed *in situ* under standard conditions; however, additional cyclic heat-induced antigen retrieval revealed distinct honeycomb-like neuronal SMARCC2 cytobodies that were increased in PD. These unique cytobodies were subsequently characterised using super-resolution microscopy in an independent study [41].

### Analysis of BCL-XL expression across multiple brain regions

TMAs enable efficient, high-throughput comparison across numerous cases but analyse only a subsection of larger tissue regions. To complement this and capture regional complexity, BCL-XL, identified as differentially expressed in both *in vitro* and *in situ* analyses, was selected for further validation in whole-tissue sections from the medulla, substantia nigra, MTG, middle frontal gyrus, and cerebellum to examine its relationship with PD progression across the brain.

BCL-XL staining was observed across all examined brain regions, consistent with TMA findings, with no observable differences in labelling between neurologically normal and PD tissues (Figure 4A-E, Supplementary Figure 5-6). Vascular BCL-XL co-localised with the pericyte marker PDGFRβ and the smooth muscle marker αSMA, showing the strongest labelling in vessels positive for both markers, while smaller PDGFRβ-only vessels displayed weaker signals. Beyond vascular localisation, BCL-XL was also detected in microglia across all regions, as indicated by co-labelling with IBA1 (Figure 4F). No quantitative regional differences were observed between PD and neurologically normal groups in the medulla, middle frontal gyrus, or cerebellum (Supplementary Table 8). In contrast, analysis of MTG whole sections confirmed the TMA results, showing higher pericytic BCL-XL levels in PD (*1*.*50-fold; 22*.*6 (nPD =11) vs 15*.*1 (nCtrl = 9); p= 0*.*034*). In the substantia nigra, BCL-XL was similarly elevated in microglia (*1*.*78-fold; 10*.*5 (nPD =11) vs 5*.*9 (nCtrl =11); p = 0*.*018*) and pericytes (*1*.*52-fold; 26*.*1 vs 17*.*2; p = 0*.*018, nPD = 11 vs nCtrl =10*) relative to neurologically normal cases (Figure 4G-K).

**Figure 4.**
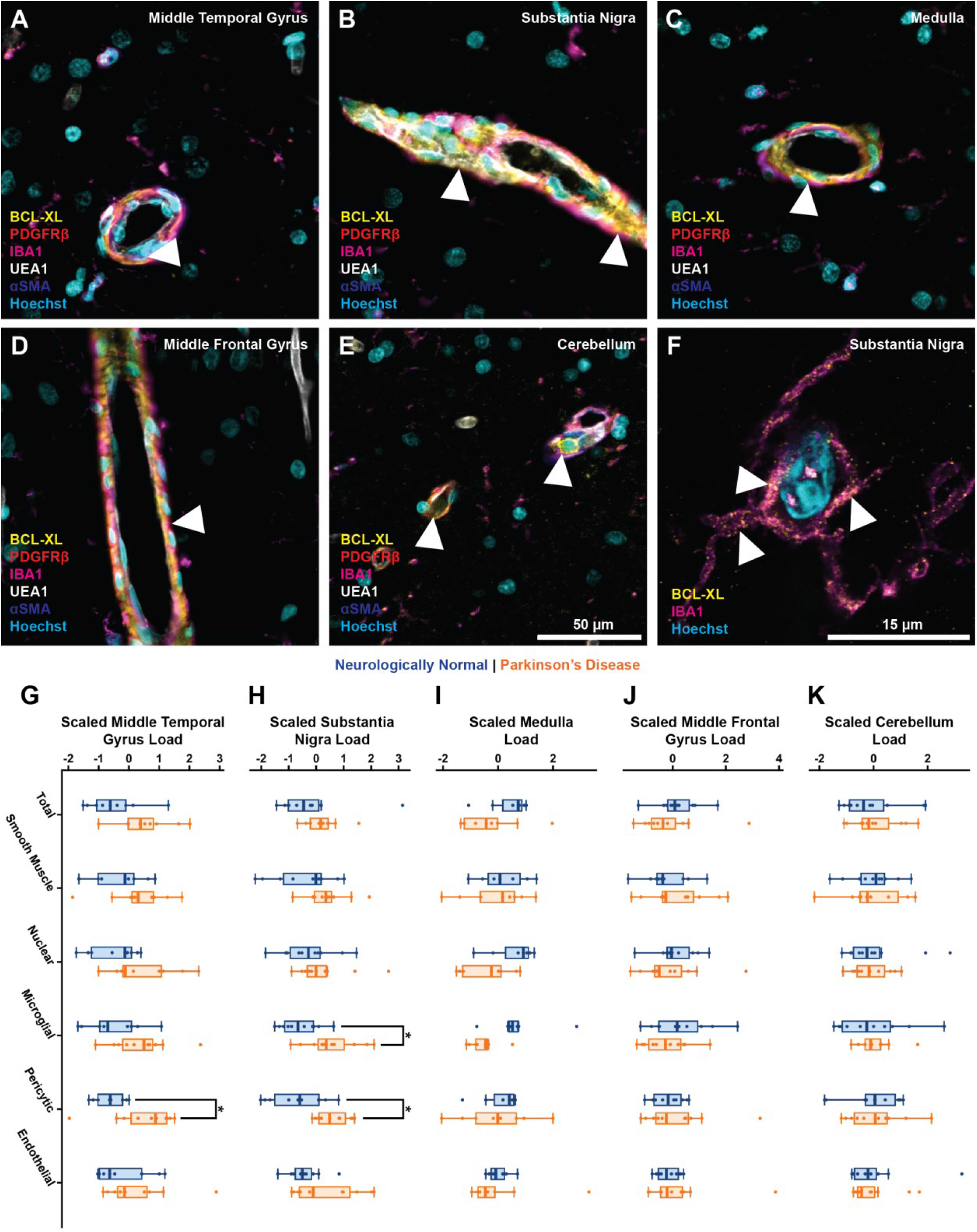
Comparative Analysis of BCL-XL expression across multiple brain regions. **(A-E)**. BCL-XL in postmortem human PD tissue across different brain regions **(F)**. Confocal Z-stack max projection image of BCL-XL staining co-localised to microglia marker IBA1 in the PD substantia nigra. White arrows indicate specific BCL-XL labelling. (G). Pericytic BCL-XL expression was increased in the PD MTG (1.50-fold, *nPD = 11 vs nCtrl = 9, p = 0*.*034*). (H). In the PD substantia nigra, BCL-XL expression was increased in microglia (1.78-fold, p = 0.018) and in pericytes (1.52-fold, nPD = 11 vs nCtrl = 10, p = 0.018). (I-K). No differences in BCL-XL expression were observed in the medulla, middle frontal gyrus, or cerebellum between neurologically normal and PD cases. Dataset normality was tested using the Shapiro-Wilk test. As appropriate, analysis was performed using a t-test or a Mann-Whitney U test, with multiple-comparison correction using the Benjamini-Hochberg method (false discovery rate = 0.05). Load (protein area normalised to tissue area) datasets were scaled using the scale function in R to preserve relative differences when creating boxplots. p < 0.05 (*) and p < 0.01 (**).

### BCL-XL expression is not associated with *α*-Syn aggregate pathology

In the MTG, BCL-XL expression showed no clear association with α-Syn labelling. Consistent with previous observations, BCL-XL staining was most prominent around blood vessels. While some vessels contained small aggregates positive for both pS129 and N-terminal α-Syn labelling, these aggregates did not appear to be directly associated with BCL-XL expression (Supplementary Figure 7). Despite the increased presence of vascular aggregates, staining patterns were similar in the SN, (Supplementary Figure 8). BCL-XL showed no consistent relationship with larger aggregates or classical Lewy bodies. Correlative analysis supported these findings, revealing no significant associations between BCL-XL expression and α-Syn load across the examined brain regions (Supplementary Table 9).

## Discussion

The search for therapeutic targets that modify α-Syn pathology remains central to synucleinopathy research. Using a strain-specific approach in primary human brain pericytes, we identified three proteins whose expression changed with α-Syn exposure: decreased NUCKS1, increased CSNK1D, and consistently elevated BCL-XL. The workflow integrated RNA-seq discovery with multi-level protein validation, linking transcriptomic signatures to stable protein changes in both cultured primary human brain-derived pericytes and post-mortem tissue. This cross-platform design enhances biological relevance and minimises the likelihood that observed differences reflect transient transcriptional noise rather than sustained cellular adaptation.

Exposure to five pure α-Syn strains (Fibrils, Fibrils-65, Fibrils-91, Fibrils-110, and Ribbons) produced 300 DEGs, revealing shared and strain-specific transcriptional patterns. This diversity mirrors known structural and biochemical variation among α-Syn fibril polymorphs [16,17,42–45] and supports the concept that distinct conformers impose discrete molecular pressures on recipient cells [46,47]. Common DEGs likely represent conserved stress or degradation pathways, whereas strain-specific DEGs may underlie heterogeneity in disease progression and regional vulnerability.

Protein analysis revealed upregulation of CSNK1D and BCL-XL at the protein level, despite decreased RNA expression, highlighting the complexity of multi-layered post-transcriptional regulation characteristic of neurodegenerative contexts [48–54]. While the short 24-hour exposure window may limit protein detection, the consistent increase across cases and the alignment of BCL-XL upregulation across methods underscore the robustness of these findings. Few studies have combined α-Syn strain–resolved transcriptomics with systematic cell-type–specific protein validation in primary human cells, providing additional strength to this integrated approach [55,56].

Among the validated proteins, BCL-XL emerged as the most consistent and biologically meaningful target. *In vitro*, α-Syn strain exposure increased BCL-XL expression in pericytes, while post-mortem analysis revealed elevated pericytic BCL-XL in the MTG and increased pericytic and microglial expression in the substantia nigra. The parallel increase in pericytes and microglia suggests coordinated stress responses across vascular and immune compartments of the neurovascular unit, consistent with evidence that pericytes actively regulate inflammatory signalling and microglial activation in neurodegenerative conditions [57]. BCL-XL, a key anti-apoptotic effector of the BCL-2 family, preserves mitochondrial integrity by neutralising BAX and BAK and preventing caspase activation [58]. Elevated BCL-XL mRNA has been reported in dopaminergic neurons in PD [59], but protein-level confirmation and cell-type specificity have not been demonstrated thus far. Our findings extend these observations to the neurovascular unit, positioning BCL-XL as a regulator of stress resilience beyond neurons.

α-Syn heterogeneity is well established across synucleinopathies, with distinct conformers linked to differential seeding capacity, regional distribution, and clinicopathological phenotype [10–12,16–18,20,60]. Modelling multiple recombinant strains enabled controlled interrogation of conformer-responsive pathways while avoiding the intrinsic variability of human tissue. In heterogeneous disease contexts where mixed-strain populations and co-pathologies coexist, such controlled systems provide a tractable platform for identifying strain-sensitive targets. Crucially, alignment of strain-identified targets with protein-level alterations in human post-mortem tissue anchors these findings in disease biology and bridges experimental and pathological contexts. Demonstrating that strain-responsive pathways, including BCL-XL upregulation, are recapitulated within the neurovascular unit strengthens translational relevance and provides a strategy for identifying vascular targets that may contribute to synucleinopathy progression.

Although recombinant α-Syn assemblies do not fully replicate the structural and post-translational complexity of human-derived aggregates, their controlled use enabled identification of conformer-responsive pathways under defined conditions. The integration of transcriptomic discovery and *in situ* validation demonstrates that pericytes translate α-Syn conformational diversity into measurable protein expression changes. Within this framework, BCL-XL emerges as a possible target at the neurovascular interface, linking pericyte responses to α-Syn pathology with adaptive stress signalling that may shape neurovascular dysfunction in Parkinson’s disease.

## Data availability

## Acknowledgements

We want to acknowledge the generosity of the brain donors and their families for their generous gift of brain tissue for research to the Neurological Foundation Human Brain Bank (Auckland, New Zealand). We also thank Marika Eszes at the Neurological Foundation Human Brain Bank and all technical staff involved in collecting and processing the human brain tissue at the Centre for Brain Research. Confocal and ImageXpress microscopy was performed using instruments in the Biomedical Imaging Research Unit at The University of Auckland. Fluorescent slide scanning was performed in Sydney. We acknowledge the technical and scientific assistance of Sydney Microscopy & Microanalysis, the University of Sydney node of Microscopy Australia. Several monoclonal antibodies (CPTC-BCL2L1-1, AFFN-FYN-7H11, PCRP-PUM2-1B11) were obtained from the Developmental Studies Hybridoma Bank, created by the NICHD of the NIH and maintained at the University of Iowa, Department of Biology, Iowa City. CPTC-BCL2L1-1 was deposited to the DSHB by Clinical Proteomics Technologies for Cancer (DSHB Hybridoma Product CPTC-BCL2L1-1). AFFN-FYN-7H11 was deposited to the DSHB by EU Program Affinomics (DSHB Hybridoma Product AFFN-FYN-7H11). PCRP-PUM2-1B11 was deposited to the DSHB by Common Fund – Protein Capture Reagents Program (DSHB Hybridoma Product PCRP-PUM2-1B11)

## Author contributions

KR, BVD, MD, and GMH contributed to the conception and design of the experiments. Experiments were performed by KR, EY and BVD. Recombinant α-Syn fibrils were prepared by RM. Human tissue and laboratory equipment were provided by GMH, RLMF, MAC, MD and BVD. Data collection and analysis were performed by KR, TS, XHW, and BR. The manuscript and figures were prepared by KR and reviewed and edited by BVD and GMH. All authors read and approved the final manuscript.

