## Supplementary Data for "Systematic screening identifies elevated neurovascular BCL-XL in Parkinson’s disease"

### Supplementary Table 1. Tissue cases used in this study

| Case | Age (Years) | Sex | PMD (Hours) | Brain Weight (g) | Cause of death | Pathology | Duration of PD (Years) |
|---|---|---|---|---|---|---|---|
| H215 | 67 | F | 23.5 | 1232 | Ischaemic heart disease | control | - |
| H180 | 73 | M | 33 | 1318 | Ischaemic heart disease | control | - |
| H184 | 35 | M | 20 | 1594 | Electrocution | control | - |
| H186 | 68 | M | 21 | 1327 | Ischaemic heart disease | control | - |
| H187 | 98 | F | 15 | 1114 | Caecal carcinoma | control | - |
| **H189**[a] | 41 | M | 16 | 1412 | Asphyxia | control | - |
| H190 | 72 | F | 19 | 1264 | Ruptured myocardial infarction | control | - |
| H194 | 68 | M | 22.5 | 1403 | Coronary atherosclerosis | control | - |
| H204 | 66 | M | 9 | 1461 | Ischaemic heart disease | control | - |
| **H209**[a] | 48 | M | 23 | 1470 | Ischaemic heart disease | control | - |
| H227 | 78 | F | 4 | | cerebrovascular accident | control | - |
| H228 | 87 | F | 21 | | Ischaemic heart disease | control | - |
| H231 | 65 | M | 8 | 1527 | Ischaemic heart disease | control | - |
| H237 | 81 | M | 17 | | acute bacterial endocarditis | control | - |
| H238 | 63 | F | 16 | 1324 | Dissecting aortic aneurysm | control | - |
| **H239**[a] | 64 | M | 15.5 | 1529 | Ischemic Heart Disease | control | - |
| H242 | 61 | M | 19.5 | 1466 | Coronary atherosclerosis | control | - |
| **H243**[a] | 77 | F | 13 | 1184 | Ischaemic heart disease- coronary atherosclerosis | control | - |
| H244 | 76 | M | 16 | 1508 | Ischaemic heart disease- coronary atherosclerosis | control | - |
| **H245**[a] | 63 | M | 20 | 1194 | Asphyxia | control | - |
| H246 | 89 | M | 17 | 1130.7 | Type II myocardial infarction | control | - |
| H247 | 51 | M | 31 | 1671 | Bilateral pulmonary thromboembolism secondary to right calf deep vein thrombosis | control | - |
| H248 | 98 | M | 10.5 | - | Congestive heart failure | control | - |
| H249 | 77 | F | 17 | 1344 | B cell lymphoma / follicular | control | - |
| **H250**[a] | 93 | F | 19 | 1143.3 | Acute coronary syndrome; pneumonia | control | - |
| H252 | 94 | F | 13.5 | 981.8 | Lower respiratory tract infection | control | - |
| H254 | 87 | M | 26 | 1440 | Cardiac arrhythmia | control | - |
| PD18 | 98 | F | 14 | 880 | Bronchopneumonia | PD | |
| PD30 | 82 | M | 19 | 1222 | Multiple organ failure | PD/CLBD | - |
| PD31 | 67 | M | 25 | 1405 | Respiratory failure | PD | 5 |
| PD32 | 71 | M | 8 | 1490 | Ischaemic heart disease/ congestive heart failure | PD/CLBD | 25 |
| PD33 | 91 | M | 4 | 1201 | Pneumonia | PD/CLBD/AD | |
| PD35 | 73 | M | 16 | 1293 | Pneumonia | PD/CLBD/ mild AD | 15 |
| PD36 | 78 | F | 22.5 | 1113 | Bronchopneumonia- CVD | PD | - |
| PD37 | 81 | M | 4 | 1190 | PD | PD/CLBD | 13 |
| PD41 | 81 | M | 13 | 1248 | Renal failure/urinary sepsis | PD/AD | 1 |
| PD43 | 60 | F | 15.5 | 1118 | Bronchopneumonia & multisystem organ failure | CLBD | 7 |
| PD50 | 88 | M | 6 | 1218 | Myocardial infarction/ ischaemic heart disease | PD/CLBD | 10 |
| **PD52**[a] | 84 | M | 5 | 1067 | Acute myocardial infarction | PD/CLBD | 12 |
| PD56 | 74 | M | 10.5 | 1322 | End-stage Lewy body disease | PD/CLBD | 12 |
| PD57 | 90 | F | 14 | 1134 | Bronchopneumonia | CLBD/AD | - |
| PD60 | 80 | M | 18 | | Urosepsis | PD/CLBD/(low AD) | - |
| **PD63**[a] | 91 | F | 5 | 1119.6 | Parkinson's disease | PD/mild AD /CLBD/ | 22 |
| **PD65**[a] | 67 | M | 2.25 | 1033.1 | Advanced PD and LBD | PD/CLBD | 9 |
| PD66 | 73 | M | 17.5 | 1376.4 | Aspiration pneumonia | PD/CLBD | 22 |
| PD67 | 65 | M | 17 | 1223.7 | Bronchopneumonia | PD/CLBD | 12 |
| PD68 | 70 | F | 18 | 989 | Parkinson's Disease | PD | |
| PD69 | 84 | F | 22 | | deconditioning, PD | PD/CLBD | - |
| **PD71**[a] | 80 | M | 5.5 | | Pneumonia | PD/CLBD | 9 |
| PD73 | 83 | F | 4 | | Pneumonia/PD | PD/LBD | - |
| **PD77**[a] | 76 | F | 6.5 | 1074.3 | Intra-abdominal high-grade serous carcinoma | PD/LBD | 23 |
| **PD78**[a] | 80 | M | 5.5 | | PD | PD/CLBD | - |
| PD79 | 77 | M | 6.5 | 1279.5 | Severe LB dementia, cognitive loss, Loss of oral intake | CLBD/PD | 22 |
| PD86 | 69 | F | 5 | 1224 | Bronchopneumonia | PD/CLBD | |
| PD90 | 77 | M | 28.75 | 1333 | Influenza A, Heart failure | PD/CLBD | |
| PD96 | 95 | M | 4.75 | 1249 | Aspiration pneumonia | PD | |
| PD105 | 82 | M | 20 | 1364.9 | Lewy Body Dementia | PD/CLBD | |

[a] pericytes derived from these cases were cultured and used for α-Synuclein treatment assays, RNAseq and immunocytochemistry

### Supplementary Table 2. Primary antibodies for analogous proteins of differentially expressed genes selected for validation.

| Gene | Protein | Catalogue | Antibody / Lectin | Source | Species | Subtype | Dilution | Antibody Registry RRID | Strain Association | | | | |
|---|---|---|---|---|---|---|---|---|---|---|---|---|---|
| | | | | | | | | | Fibril | Fibril-65 | Fibril-91 | Fibril-110 | Ribbon |
| ABCF1 | ATP Binding Cassette Subfamily F Member 1 | PA5-82591 | ABCF1 Polyclonal Antibody | ThermoFisher Scientific | Rabbit | IgG | 1:50 | AB_2789748 | ✓ | | | | |
| AHSA1 | Activator Of HSP90 ATPase Activity 1 | ab91259 | Anti-AHA1 antibody (25F2.D9) | Abcam | Rat | IgG2a | 1:50 | AB_2049149 | | | | | ✓ |
| ARHGEF7 | Rho Guanine Nucleotide Exchange Factor 7 | sc-393184 | βPIX Antibody (H-3) | Santa Cruz | Mouse | IgG1 | 1:50 | AB_3096150 | | ✓ | | | |
| ASAH1 | N-Acylsphingosine Amidohydrolase 1 | sc-136275 | Acid Ceramidase Antibody (23) | Santa Cruz | Mouse | IgG1 | 1:50 | AB_10609901 | | | | | ✓ |
| ATP6V1D | ATPase H+ Transporting V1 Subunit D | sc-390384 | V-ATPase D Antibody (E-12) | Santa Cruz | Mouse | IgG1 | 1:50 | AB_3096158 | | | | ✓ | |
| BCLXL | BCL2 Like 1 | CPTC-BCL2L1-1 | CPTC-BCL2L1-1 | DSHB | Mouse | IgG2c | 1:50 | AB_10805150 | | | | | |
| CD226 | CD226 Molecule | sc-376736 | DNAM-1 Antibody (D-11) | Santa Cruz | Mouse | IgG1 | 1:50 | AB_3096147 | ✓ | ✓ | | | ✓ |
| CSNK1D | Casein Kinase 1 Delta | sc-55553 | casein kinase Iδ Antibody (C-8) | Santa Cruz | Mouse | IgG2a | 1:50 | AB_831049 | | | | ✓ | |
| CSNK2A1 | Casein Kinase 2 Alpha 1 | AFFN-CSNK2A1-3E6 | AFFN-CSNK2A1-3E6 | DSHB | Mouse | IgG2b | 1:50 | AB_2721968 | | | | | ✓ |
| CSNK2B | Casein Kinase 2 Beta | sc-12739 | casein kinase IIβ Antibody (6D5) | Santa Cruz | Mouse | IgG1 | 1:50 | AB_62679) | ✓ | | | ✓ | |
| EBAG9 | Estrogen Receptor Binding Site Associated Antigen 9 | CPTC-EBAG9-1 | CPTC-EBAG9-1 | DSHB | Mouse | IgG2c | 1:50 | AB_2617249 | ✓ | | | | ✓ |
| EIF4E2 | Eukaryotic Translation Initiation Factor 4E Family Member 2 | sc-100731 | eIF4E2 Antibody (YB-18) | Santa Cruz | Mouse | IgG1 | 1:50 | AB_1122491 | | | ✓ | | |
| ELF4 | E74 Like ETS Transcription Factor 4 | sc-515363 | Elf-4 Antibody (E-11) | Santa Cruz | Mouse | IgM | 1:50 | AB_3096151 | ✓ | ✓ | | | ✓ |
| EML2 | EMAP Like 2 | sc-374627 | EML2 Antibody (F-3) | Santa Cruz | Mouse | IgG1 | 1:50 | AB_10986270 | | | | | |
| FYN | FYN Proto-Oncogene, Src Family Tyrosine Kinase | AFFN-FYN-7H11 | AFFN-FYN-7H11 | DSHB | Mouse | IgG2b | 1:50 | AB_261760 | | | | ✓ | |
| GEM | GTP Binding Protein Overexpressed In Skeletal Muscle | sc-166891 | Gem Antibody (G-1) | Santa Cruz | Mouse | IgG$_{2a}$ | 1:50 | AB_10610512 | | ✓ | | | ✓ |
| GNA15 | G Protein Subunit Alpha 15 | sc-393878 | Gα 15 Antibody (F-3) | Santa Cruz | Mouse | IgG1 | 1:50 | AB_2734138 | | ✓ | | ✓ | |
| ICA1 | Islet Cell Autoantigen 1 | sc-271489 | ICA69 Antibody (A-1) | Santa Cruz | Mouse | IgM | 1:50 | AB_10648479 | | ✓ | ✓ | | ✓ |
| IGFBP3 | Insulin Like Growth Factor Binding Protein 3 | sc-365936 | IGFBP3 Antibody (B-5) | Santa Cruz | Mouse | IgG2b | 1:50 | AB_10917037 | | | ✓ | | |
| INO80B | INO80 Complex Subunit B | sc-390009 | INO80B Antibody (E-3) | Santa Cruz | Mouse | IgG2b | 1:50 | AB_3096155 | | | ✓ | | |
| ITGA3 | Integrin Subunit Alpha 3 | P1B5 | P1B5 | DSHB | Mouse | IgG1 | 1:50 | AB_2619595 | | | | | ✓ |
| KIF21B | Kinesin Family Member 21B | HPA027274 | Anti-KIF21B antibody | Sigma Aldrich | Rabbit | IgG | 1:50 | AB_1852483 | ✓ | | | | ✓ |
| KLF5 | KLF Transcription Factor 5 | PCRP-KLF5-1A3 | PCRP-KLF5-1A3 | DSHB | Mouse | IgG2b | 1:50 | AB_2618808 | | | | | |
| KRAS | KRAS Proto-Oncogene, GTPase | CPTC-KRAS4B-1 | CPTC-KRAS4B-1 | DSHB | Mouse | IgG1 | 1:50 | AB_2722074 | | ✓ | | | |
| LRRC10B | Leucine Rich Repeat Containing 10B | ab188260 | Anti-LRRC10B antibody | Abcam | Rabbit | IgG1 | 1:50 | AB_3096148 | ✓ | ✓ | ✓ | ✓ | ✓ |
| MAT2B | Methionine Adenosyltransferase 2 Non-Catalytic Beta Subunit | PCRP-MAT2B-1G7 | PCRP-MAT2B-1G7 | DSHB | Mouse | IgG2b | 1:50 | AB_2618831 | ✓ | | | | ✓ |
| MED22 | Mediator Complex Subunit 22 | PCRP-MED22-1E4 | PCRP-MED22-1E4 | DSHB | Mouse | IgG2c | 1:50 | AB_2722241 | | | | ✓ | |
| MEGF11 | Multiple EGF-like domains protein 11 | PA5107171 | MEGF11 Polyclonal Antibody | ThermoFisher Scientific | Rabbit | IgG | 1:2000 | AB_2817887 | | | ✓ | | ✓ |
| MTHFD1 | Methylenetetrahydrofolate Dehydrogenase, Cyclohydrolase And Formyltetrahydrofolate Synthetase 1 | sc-376722 | MTHFD1/1L Antibody (D-9) | Santa Cruz | Mouse | IgG2b | 1:50 | AB_3096149 | | ✓ | ✓ | ✓ | ✓ |
| NR3C1 | Nuclear Receptor Subfamily 3 Group C Member 1 | sc-393232 | GR/NR3C1/Glucocorticoid Receptor Antibody (F-10) | Santa Cruz | Mouse | IgG2b | 1:50 | AB_2687823 | ✓ | | | | |
| NUB1 | Negative Regulator Of Ubiquitin Like Proteins 1 | sc-377003 | NUB1 Antibody (F-10) | Santa Cruz | Mouse | IgG1 | 1:50 | AB_3096146 | ✓ | ✓ | | | |
| NUCKS1 | Nuclear Casein Kinase And Cyclin Dependent Kinase Substrate 1 | HPA062351 | Anti-NUCKS1 antibody produced in rabbit | Sigma Aldrich | Rabbit | IgG | 1:1000 | AB_2684745 | | | | | ✓ |
| P3H2 | Prolyl 3-Hydroxylase 2 | sc-377096 | LEPREL1 Antibody (H-4) | Santa Cruz | Mouse | IgG1 | 1:50 | AB_3096156 | | | ✓ | | ✓ |
| PHGDH | Phosphoglycerate Dehydrogenase | AFFN-PHGDH-13A8-B1 | AFFN-PHGDH-13A8-B1 | DSHB | Mouse | IgG1 | 1:50 | AB_2617819 | ✓ | | | | |
| PUM2 | Pumilio RNA Binding Family Member 2 | PCRP-PUM2-1B11 | PCRP-PUM2-1B11 | DSHB | Mouse | IgG2c | 1:50 | AB_2722308 | ✓ | | | | |
| RXRA | Retinoid X Receptor Alpha | PCRP-RXRA-2A8 | PCRP-RXRA-2A8 | DSHB | Mouse | IgG2a | 1:50 | AB_2619055 | | | | ✓ | |
| SMARCC2 | SWI/SNF Related, Matrix Associated, Actin Dependent Regulator Of Chromatin Subfamily C Member 2 | sc17838 | BAF170 Antibody (E-6) | Santa Cruz | Mouse | IgG1 | 1:50 | AB_2286337 | | | | ✓ | |
| TAPBP | TAP Binding Protein | CPTC-TAPBP-1 | CPTC-TAPBP-1 | DSHB | Mouse | IgG2c | 1:50 | AB_2814788 | | | ✓ | | |
| TAX1BP1 | Tax1 Binding Protein 1 | PCRP-TAX1BP1-1D4 | PCRP-TAX1BP1-1D4 | DSHB | Mouse | IgG2a | 1:50 | AB_2619161 | | | | | ✓ |
| TGIF1 | TGFB Induced Factor Homeobox 1 | PCRP-TGIF1-2G6 | PCRP-TGIF1-2G6 | DSHB | Mouse | IgG1 | 1:50 | AB_2619189 | | | | | ✓ |
| ZFYVE19 | Zinc Finger FYVE-Type Containing 19 | PCRP-ZFYVE19-3C5 | PCRP-ZFYVE19-3C5 | DSHB | Mouse | IgG2b | 1:50 | AB_2619287 | | | ✓ | | |
| ZHX2 | Zinc Fingers And Homeoboxes 2 | PCRP-ZHX2-1B2 | PCRP-ZHX2-1B2 | DSHB | Mouse | IgG2a | 1:50 | AB_2619293 | | ✓ | | | |
| ZNF24 | Zinc Finger Protein 24 | PCRP-ZNF24-1E12 | PCRP-ZNF24-1E12 | DSHB | Mouse | IgG2a | 1:50 | AB_2619361 | | ✓ | ✓ | ✓ | |

DSHB – Developmental Studies Hybridoma Bank

### Supplementary Table 3. Primary cell-type-specific antibodies and lectins

| Gene | Cell Type | Protein | Catalogue# | Antibody / Lectin | Source | Species | Subtype | Dilution | Antibody Registry RRID |
|---|---|---|---|---|---|---|---|---|---|
| AQP4 | Astrocytes | Aquaporin 4 | AB3594 | Anti-Aquaporin 4 Antibody, CT | Sigma Aldrich | Rabbit | IgG | 1:800 | AB_91530 |
| BCL2L1 | - | BCL-XL | 10783-1-AP | BCL2L1 Polyclonal antibody | ProteinTech | Rabbit | IgG | 1:500 | AB_2064724 |
| GFAP | Astrocytes | Glial Fibrillary Acidic Protein | Ab4674 | Anti-GFAP antibody | Abcam | Chicken | IgY | 1:2000 | AB_304558 |
| - | Endothelial Cells | - | L8262 | Lectin from Ulex europaeus (gorse, furze) | Sigma Aldrich | - | - | 1:1000 | - |
| IBA1 | Microglia | Allograft Inflammatory Factor 1 | 234009 | Anti-IBA1 antibody - 234 009 | Synaptic Systems | Chicken | IgY | 1:1000 | AB_2891282 |
| NeuN | Neurons | RNA Binding Fox-1 Homolog 3 | ABN90P | Anti-NeuN purified Antibody | Sigma Aldrich | Guinea Pig | IgG | 1:500 | AB_2341095 |
| PDGFRβ | Pericytes | Platelet Derived Growth Factor Receptor Beta | RDSAF385 | Anti-Human PDGFR beta Antibody | R&D Systems | Goat | IgG | 1:500 | AB_355339 |
| ACTA2 | Smooth Muscle Cells | Actin Alpha 2, Smooth Muscle | ab5694 | Anti-alpha smooth muscle Actin antibody | Abcam | Rabbit | IgG | 1:200 | AB_2223021 |
| SNCA | - | α-Synuclein (N-Terminus Labelling) | 849101 | Purified anti-α-Synuclein, 34-45 Antibody | BioLegend | Mouse | IgG1 | 1:1000 | AB_2650703 |
| SNCA | | α-Synuclein (PS129 Labelling) | ab184674 | Anti-Alpha-synuclein (phospho S129) antibody [P-syn/81A] | Abcam | Mouse | IgG2a | 1:4000 | AB_2819037 |

### Supplementary Table 4. Secondary antibodies, fluorophores and nuclear dyes

| Target | Catalogue# | Antibody | Source | Species | Dilution | Antibody Registry RRID |
|---|---|---|---|---|---|---|
| Mouse | A-21202 | Donkey anti-Mouse IgG (H+L) Highly Cross-Adsorbed Secondary Antibody, Alexa Fluor™ 488 | ThermoFisher Scientific | Donkey | 1:500 | AB_141607 |
| Mouse | smsG1CL488-1 | ChromoTek Nano-Secondary® alpaca anti-mouse IgG1, recombinant VHH, CoraLite® Plus 488 (CTK0103, CTK0104) | Proteintech | Alpaca | 1:500 | AB_2941310 |
| Mouse | A-11029 | Goat anti-Mouse IgG (H+L) Highly Cross-Adsorbed Secondary Antibody, Alexa Fluor™ 488 | ThermoFisher Scientific | Goat | 1:500 | AB_2534088 |
| Mouse | 115-547-188 | Alexa Fluor® 488 AffiniPure™ Fab Fragment Goat Anti-Mouse IgG2c, Fcγ fragment specific | Jackson Immuno-Research | Goat | 1:500 | AB_2632537 |
| Rabbit | A-21206 | Donkey anti-Rabbit IgG (H+L) Highly Cross-Adsorbed Secondary Antibody, Alexa Fluor™ 488 | Thermo Fisher Scientific | Donkey | 1:500 | AB_2535792 |
| Rabbit | A-11034 | Goat anti-Rabbit IgG (H+L) Highly Cross-Adsorbed Secondary Antibody, Alexa Fluor™ 488 | ThermoFisher Scientific | Goat | 1:500 | AB_2576217 |
| Rabbit | A-11035 | Goat anti-Rabbit IgG (H+L) Highly Cross-Adsorbed Secondary Antibody, Alexa Fluor™ 546 | ThermoFisher Scientific | Donkey | 1:500 | AB_2534093 |
| Rabbit | A-31572 | Donkey anti-Rabbit IgG (H+L) Highly Cross-Adsorbed Secondary Antibody, Alexa Fluor™ 555 | ThermoFisher Scientific | Donkey | 1:500 | AB_162543 |
| Chicken | A78949 | Donkey anti-Chicken IgY (H+L) Highly Cross Adsorbed Secondary Antibody, Alexa Fluor™ 555 | ThermoFisher Scientific | Donkey | 1:500 | AB_2921071 |
| Rabbit | A-11011 | Goat anti-Rabbit IgG (H+L) Cross-Adsorbed Secondary Antibody, Alexa Fluor™ 568 | ThermoFisher Scientific | Goat | 1:500 | AB_143157 |
| Rabbit | A-11012 | Goat anti-Rabbit IgG (H+L) Cross-Adsorbed Secondary Antibody, Alexa Fluor™ 594 | ThermoFisher Scientific | Goat | 1:500 | AB_2534079 |
| Mouse | A-11032 | Goat anti-Mouse IgG (H+L) Highly Cross-Adsorbed Secondary Antibody, Alexa Fluor™ 594 | ThermoFisher Scientific | Goat | 1:500 | AB_2534091 |
| Mouse | A-21125 | Goat anti-Mouse IgG1 Cross-Adsorbed Secondary Antibody, Alexa Fluor™ 594 | ThermoFisher Scientific | Goat | 1:500 | AB_2535767 |
| Goat | A-32758 | Donkey anti-Goat IgG (H+L) Highly Cross-Adsorbed Secondary Antibody, Alexa Fluor™ Plus 594 | ThermoFisher Scientific | Donkey | 1:500 | AB_2762828 |
| Guinea Pig | 706-585-148 | Alexa Fluor® 594 AffiniPure™ Donkey Anti-Guinea Pig IgG (H+L) | Jackson Immuno-Research | Donkey | 1:500 | AB_2340474 |
| Guinea Pig | A-11076 | Goat anti-Guinea Pig IgG (H+L) Highly Cross-Adsorbed Secondary Antibody, Alexa Fluor™ 594 | ThermoFisher Scientific | Goat | 1:500 | AB_2534120 |
| Guinea Pig | 706-605-148 | Alexa Fluor® 647 AffiniPure™ Donkey Anti-Guinea Pig IgG (H+L) | Jackson Immuno-Research | Donkey | 1:500 | AB_2340476 |
| Rabbit | A-31573 | Donkey anti-Rabbit IgG (H+L) Highly Cross-Adsorbed Secondary Antibody, Alexa Fluor™ 647 | ThermoFisher Scientific | Donkey | 1:500 | AB_2536183 |
| Goat | A-21447 | Donkey anti-Goat IgG (H+L) Cross-Adsorbed Secondary Antibody, Alexa Fluor™ 647 | ThermoFisher Scientific | Donkey | 1:500 | AB_2535864 |
| Chicken | 703-605-155 | Alexa Fluor® 647 AffiniPure™ Donkey Anti-Chicken IgY (IgG) (H+L) | Jackson Immuno-Research | Donkey | 1:500 | AB_2340379 |
| Chicken | A-21449 | Goat anti-Chicken IgY (H+L) Secondary Antibody, Alexa Fluor™ 647 | ThermoFisher Scientific | Goat | 1:500 | AB_2535866 |
| - | 016-170-084 | Cy™5 Streptavidin | Jackson Immuno-Research | - | 1:500 | AB_2337245 |
| Mouse | 926-32352 | IRDye® 800CW Goat anti-Mouse IgG2b-Specific Secondary Antibody | Li-Cor | Goat | 1:500 | AB_2782999 |
| Chicken | 926-32218 | IRDye® 800CW Donkey anti-Chicken Secondary Antibody | Li-Cor | Donkey | 1:500 | AB_1850023 |
| - | 21851 | Streptavidin Protein, DyLight™ 800 | ThermoFisher Scientific | - | 1:500 | - |
| DNA | H3570 | Hoechst 23342 | ThermoFisher Scientific | - | 1:20000* | - |

### Supplementary Table 5. Optimised antigen retrieval methods for DEG analogous protein antibodies

| Gene | Protein | Antibody | Tris-EDTA (pH 9) | Sodium citrate (pH 6) | Tris-EDTA (pH 9) + Formic acid | Sodium citrate (pH 6) + Formic acid |
|---|---|---|---|---|---|---|
| **ABCF1** | **ATP Binding Cassette Subfamily F Member 1** | **ABCF1 Polyclonal Antibody** | ✓ | | | |
| AHSA1 | Activator Of HSP90 ATPase Activity 1 | Anti-AHA1 antibody (25F2.D9) | × | × | × | × |
| ARHGEF7 | Rho Guanine Nucleotide Exchange Factor 7 | βPIX Antibody (H-3) | × | × | × | × |
| **ASAH1** | **N-Acylsphingosine Amidohydrolase 1** | **Acid Ceramidase Antibody (23)** | ✓ | | | |
| ATP6V1D | ATPase H+ Transporting V1 Subunit D | V-ATPase D Antibody (E-12) | × | × | × | × |
| **BCLXL** | **BCL2 Like 1** | **CPTC-BCL2L1-1** | ✓ | | | |
| CD226 | CD226 Molecule | DNAM-1 Antibody (D-11) | × | × | × | × |
| CSNK1D | Casein Kinase 1 Delta | casein kinase Iδ Antibody (C-8) | × | × | × | × |
| CSNK2A1 | Casein Kinase 2 Alpha 1 | AFFN-CSNK2A1-3E6 | × | × | × | × |
| **CSNK2B** | **Casein Kinase 2 Beta** | **casein kinase IIβ Antibody (6D5)** | ✓ | | | |
| EBAG9 | Estrogen Receptor Binding Site Associated Antigen 9 | CPTC-EBAG9-1 | × | × | × | × |
| EIF4E2 | Eukaryotic Translation Initiation Factor 4E Family Member 2 | eIF4E2 Antibody (YB-18) | × | × | × | × |
| ELF4 | E74 Like ETS Transcription Factor 4 | Elf-4 Antibody (E-11) | × | × | × | × |
| EML2 | EMAP Like 2 | EML2 Antibody (F-3) | × | × | × | × |
| **FYN** | **FYN Proto-Oncogene, Src Family Tyrosine Kinase** | **AFFN-FYN-7H11** | ✓ | | | |
| **GEM** | **GTP Binding Protein Overexpressed In Skeletal Muscle** | **Gem Antibody (G-1)** | ✓ | | | |
| GNA15 | G Protein Subunit Alpha 15 | Gα 15 Antibody (F-3) | × | × | × | × |
| ICA1 | Islet Cell Autoantigen 1 | ICA69 Antibody (A-1) | × | × | × | × |
| IGFBP3 | Insulin Like Growth Factor Binding Protein 3 | IGFBP3 Antibody (B-5) | × | × | × | × |
| INO80B | INO80 Complex Subunit B | INO80B Antibody (E-3) | × | × | × | × |
| ITGA3 | Integrin Subunit Alpha 3 | P1B5 | × | × | × | × |
| KIF21B | Kinesin Family Member 21B | Anti-KIF21B antibody | × | × | × | × |
| KLF5 | KLF Transcription Factor 5 | PCRP-KLF5-1A3 | × | × | × | × |
| KRAS | KRAS Proto-Oncogene, GTPase | CPTC-KRAS4B-1 | × | × | × | × |
| LRRC10B | Leucine Rich Repeat Containing 10B | Anti-LRRC10B antibody | × | × | × | × |
| MAT2B | Methionine Adenosyltransferase 2 Non-Catalytic Beta Subunit | PCRP-MAT2B-1G7 | × | × | × | × |
| MED22 | Mediator Complex Subunit 22 | PCRP-MED22-1E4 | × | × | × | × |
| **MEGF11** | **Multiple EGF-like domains protein 11** | **MEGF11 Polyclonal Antibody** | ✓ | | | |
| **MTHFD1** | **Methylenetetrahydrofolate Dehydrogenase, Cyclohydrolase And Formyltetrahydrofolate Synthetase 1** | **MTHFD1/1L Antibody (D-9)** | ✓ | | ✓ | |
| NR3C1 | Nuclear Receptor Subfamily 3 Group C Member 1 | GR/NR3C1/Glucocorticoid Receptor Antibody (F-10) | × | × | × | × |
| NUB1 | Negative Regulator Of Ubiquitin Like Proteins 1 | NUB1 Antibody (F-10) | × | × | × | × |
| **NUCKS1** | **Nuclear Casein Kinase And Cyclin Dependent Kinase Substrate 1** | **Anti-NUCKS1 antibody produced in rabbit** | ✓ | | | |
| P3H2 | Prolyl 3-Hydroxylase 2 | LEPREL1 Antibody (H-4) | × | × | × | × |
| PHGDH | Phosphoglycerate Dehydrogenase | AFFN-PHGDH-13A8-B1 | × | × | × | × |
| **PUM2** | **Pumilio RNA Binding Family Member 2** | **PCRP-PUM2-1B11** | × | ✓ | | |
| RXRA | Retinoid X Receptor Alpha | PCRP-RXRA-2A8 | × | × | × | × |
| **SMARCC2** | **SWI/SNF Related, Matrix Associated, Actin Dependent Regulator Of Chromatin Subfamily C Member 2** | **BAF170 Antibody (E-6)** | ✓ | | | |
| TAPBP | TAP Binding Protein | CPTC-TAPBP-1 | × | × | × | × |
| TAX1BP1 | Tax1 Binding Protein 1 | PCRP-TAX1BP1-1D4 | × | × | × | × |
| TGIF1 | TGFB Induced Factor Homeobox 1 | PCRP-TGIF1-2G6 | × | × | × | × |
| ZFYVE19 | Zinc Finger FYVE-Type Containing 19 | PCRP-ZFYVE19-3C5 | × | × | × | × |
| ZHX2 | Zinc Fingers And Homeoboxes 2 | PCRP-ZHX2-1B2 | × | × | × | × |
| ZNF24 | Zinc Finger Protein 24 | PCRP-ZNF24-1E12 | × | × | × | × |

✓- with blue shading indicates optimised antigen retrieval method. Grey shading indicates additional antigen retrieval method was not attempted. × - antigen retrieval optimisation attempted but unsuccessful

### Supplementary Methods

### Additional antibody validation BCL-XL

In addition to the validation and characterisation data (antibodies.cancer.gov/detail/CPTC-BCL2L1-1#) available for the BCL-XL antibody (CPTC-BCL2L1-1, DSHB) used in this study, we performed further validation by overexpressing BCL-XL and labelling the cells with an additional antibody (#10783-1-AP, Proteintech). Our results demonstrated a clear one-to-one overlap in labelling between overexpressed BCL-XL and both antibodies, further confirming the specificity of the BCL-XL antibodies (**Supplementary Figure 1)**.

**HEK293 cells cell culture and transfection**
Commercial HEK293 cells (Microbix Biosystems Inc.) were cultured in complete DMEM:F12 medium, with half-medium changes every four days, until cultures reached approximately 90% confluency. HEK293 cells were passaged and plated using the same procedures described for pericytes. Cells were seeded in 6 and 96-well plates in complete DMEM:F12 24 hours prior to transfection to achieve approximately 80% confluency at the time of transfection.

Transfection was performed using Lipofectamine™ 3000 reagent (#L3000001, Thermo Fisher Scientific) according to the manufacturer's protocol. Briefly, BCL-XL expression plasmids (Venus-BclXL-pEGFP-C1; a gift from David Andrews, Addgene plasmid #177409; http://n2t.net/addgene:177409; RRID:Addgene_177409) were diluted in Opti-MEM™ Reduced Serum Medium (#31985062, Thermo Fisher Scientific) before adding P3000™ reagent. Separately, Lipofectamine™ 3000 reagent was diluted in Opti-MEM. The DNA–P3000 mixture was then combined with the Lipofectamine 3000 solution, gently mixed, and incubated at room temperature for 15 minutes to allow complex formation. Before transfection, the culture medium was replaced with fresh complete DMEM:F12, and the transfection complex was added dropwise to each well. Cells were incubated at 37°C in 5% $CO_2$ for 72 hours. 96 well plates were then fixed with 4% PFA for 15 minutes, while protein was harvested from 6 well plates.

**Protein extraction from cell cultures**
Conditioned media was removed from all wells, and cells were washed twice with 4˚C PBS. Cells were harvested using a cell scraper in 100µl sample harvesting buffer (62.5 mM Tris- HCl pH 6.8, 2% SDS, 10% glycerol), transferred to an Eppendorf and incubated at 100°C for 10 minutes. Samples were then centrifuged at 1000 rpm for 1 minute and stored at -80°C.

**Immunocytochemical validation**
Transfected cell cultures in 96 well plates were labelled via immunocytochemistry as described in the main text. Cultures were incubated with two BCL-XL antibodies (#CPTC-BCL2L1-1, DSHB; and #10783-1-AP, Proteintech) before labelling with the appropriate secondary antibodies. Fluorescently stained cells were imaged as detailed in the main text, and acquired images were analysed to verify co-localisation between antibody labelling and Venus-BCL-XL fluorescence.

**Protein extraction from human tissue**
Fresh frozen human tissue was homogenized in sample harvesting buffer (62.5 mM Tris-HCl pH 6.8, 2% SDS, 10% glycerol) at 100°C for 10 minutes. Tissue lysates were centrifuged at 10000 g for 10 minutes before protein supernatant was removed and stored at -80°C.

**Western blotting**

Protein extracts were denatured in NuPage LDS sample buffer (#NP0007, ThermoFisher Scientific) at 85°C for 5 minutes before being loaded into 4-12% Bis-Tris gels (#NP0336BOX, ThermoFisher Scientific) as described in Carmichael-Lowe et al. 2024. High-resolution separation of proteins through SDS-PAGE was conducted using MOPS SDS running buffer (#J62847-AP, ThermoFisher Scientific). A dual colour Protein Ladder was used in all gels (#1610374, BioRad). Gels were transferred onto methanol-activated PVDF membranes (#IPFL00005, Millipore) for 1 hour at a constant 20V, in NuPage transfer buffer (#NP0006, ThermoFisher Scientific). Membranes were blocked for 1 hour at room temperature in a 1:1 solution of Intercept® Blocking buffer (#927–60001, LI-COR) and TBS-T (TBS with 0.01% Tween 20), before being incubated with primary antibodies in blocking buffer at 4°C overnight. Membranes were washed with TBS-T (3 x 10 minutes) before being incubated with secondary antibodies diluted in blocking buffer with 0.02% SDS for 3 hours at room temperature, whilst light protected. Following incubation, membranes were washed in TBS-T (3 x 10 minutes) and once in TBS (10 minutes). Imaging of membranes was performed using a BioRad ChemiDoc™ MP Imaging system.

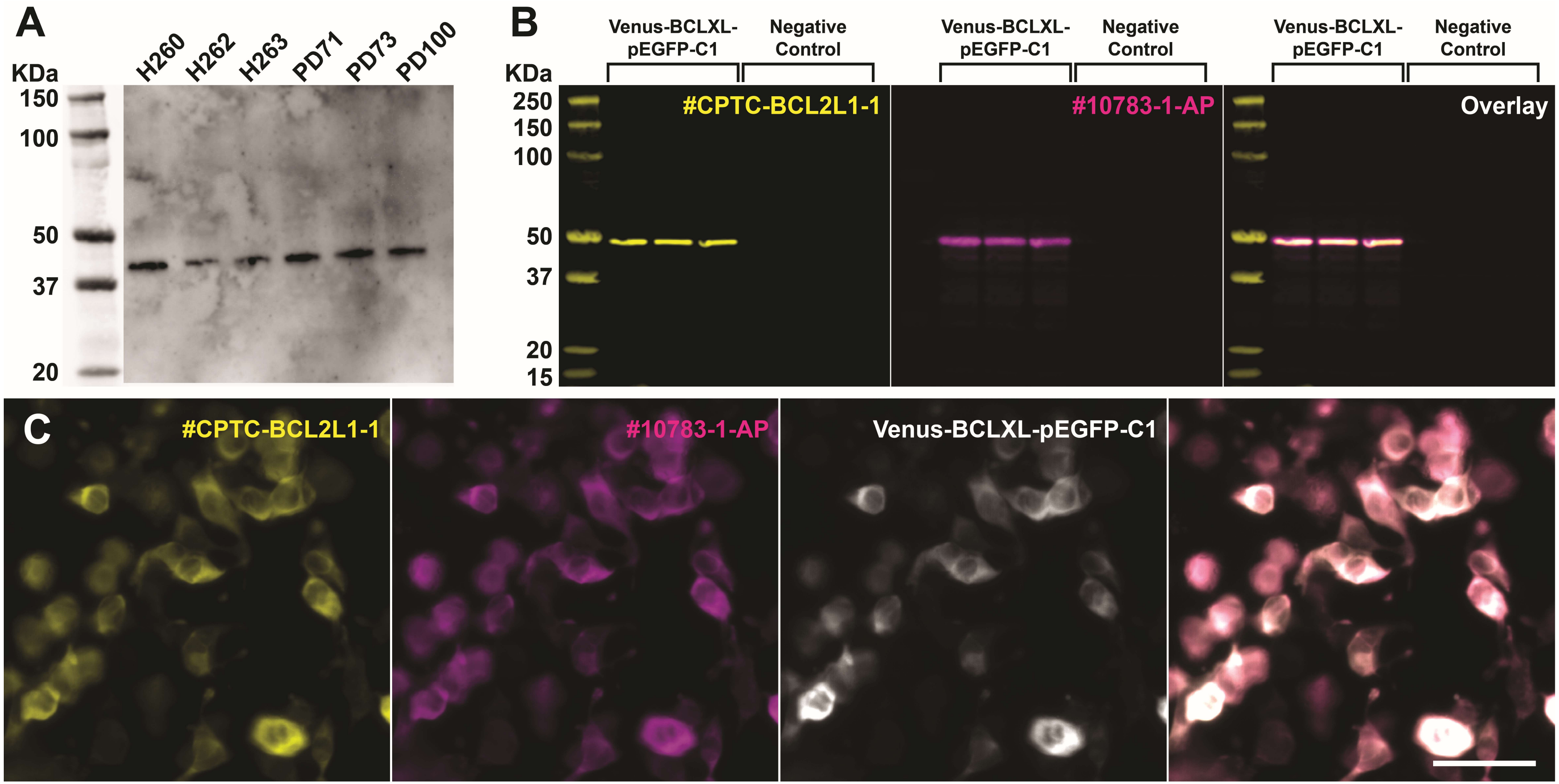


**Supplementary Figure 1. BCL-XL antibody validation.** **(A)** Western blot validation of antibody #CPTC-BCL2L1-1 using postmortem human brain tissue lysates. **(B)** Western blot validation of antibodies #CPTC-BCL2L1-1 and #10783-1-AP using Venus-BclXL-pEGFP-C1 transfected pericyte lysates. **(C)** HEK293 cells transfected with BCL-XL plasmid Venus-BclXL-pEGFP-C1were double labelled with BCL-XL antibodies #CPTC-BCL2L1-1 and #10783-1-AP to verify the specificity of #CPTC-BCL2L1-1. Overlap between labelling of both antibodies and the Venus protein indicates antibody specificity for the BCL-XL protein. Scale bar represents 50 μm.

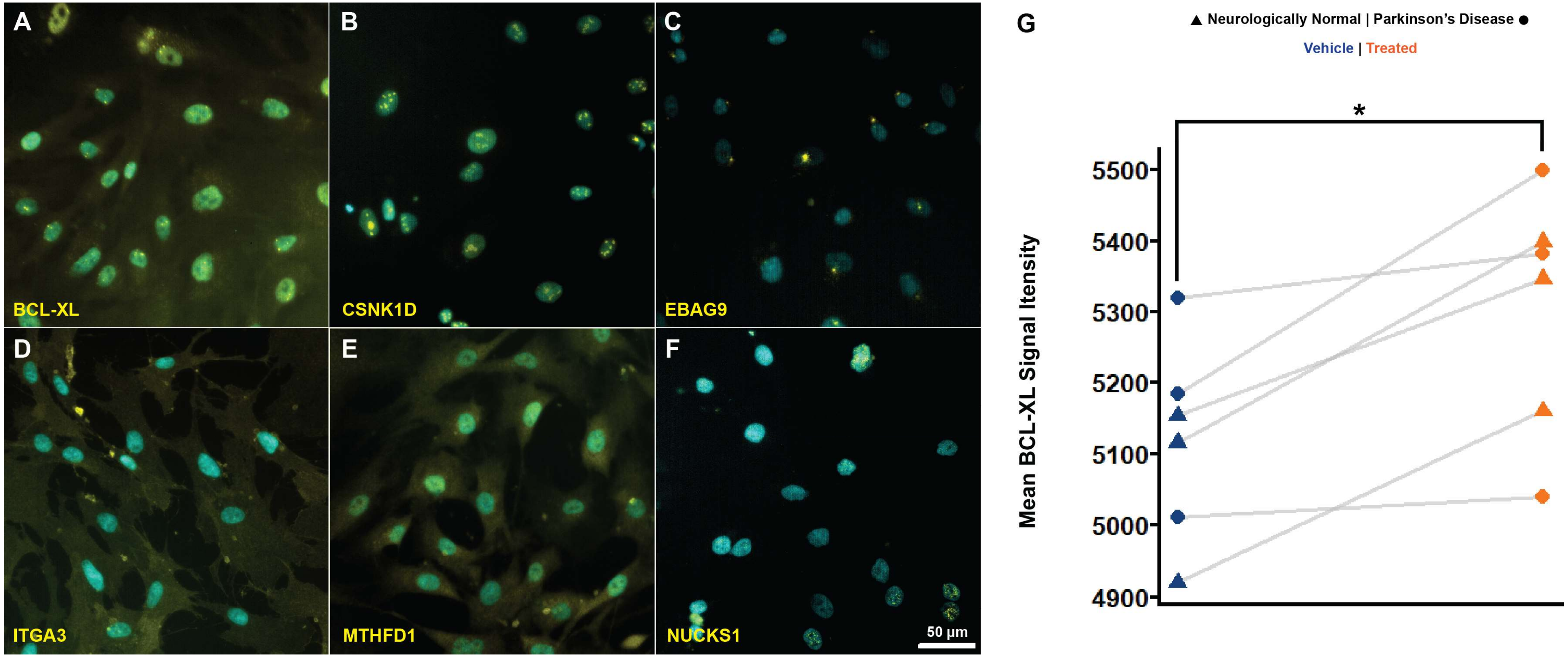


**Supplementary Figure 2. Immunocytochemical validation of differentially expressed gene analogous protein expression in vehicle treated pericytes. (A-F)** Representative images of immunocytochemical labelled proteins in vehicle treated pericytes. **(G)** Confirmation of BCL-XL protein expression via an independent experiment confirming a significant 3.7% increase in mean signal intensity in Fibril-65-treated pericytes ($p = 0.031$, Cohen's $d = 1.59$).

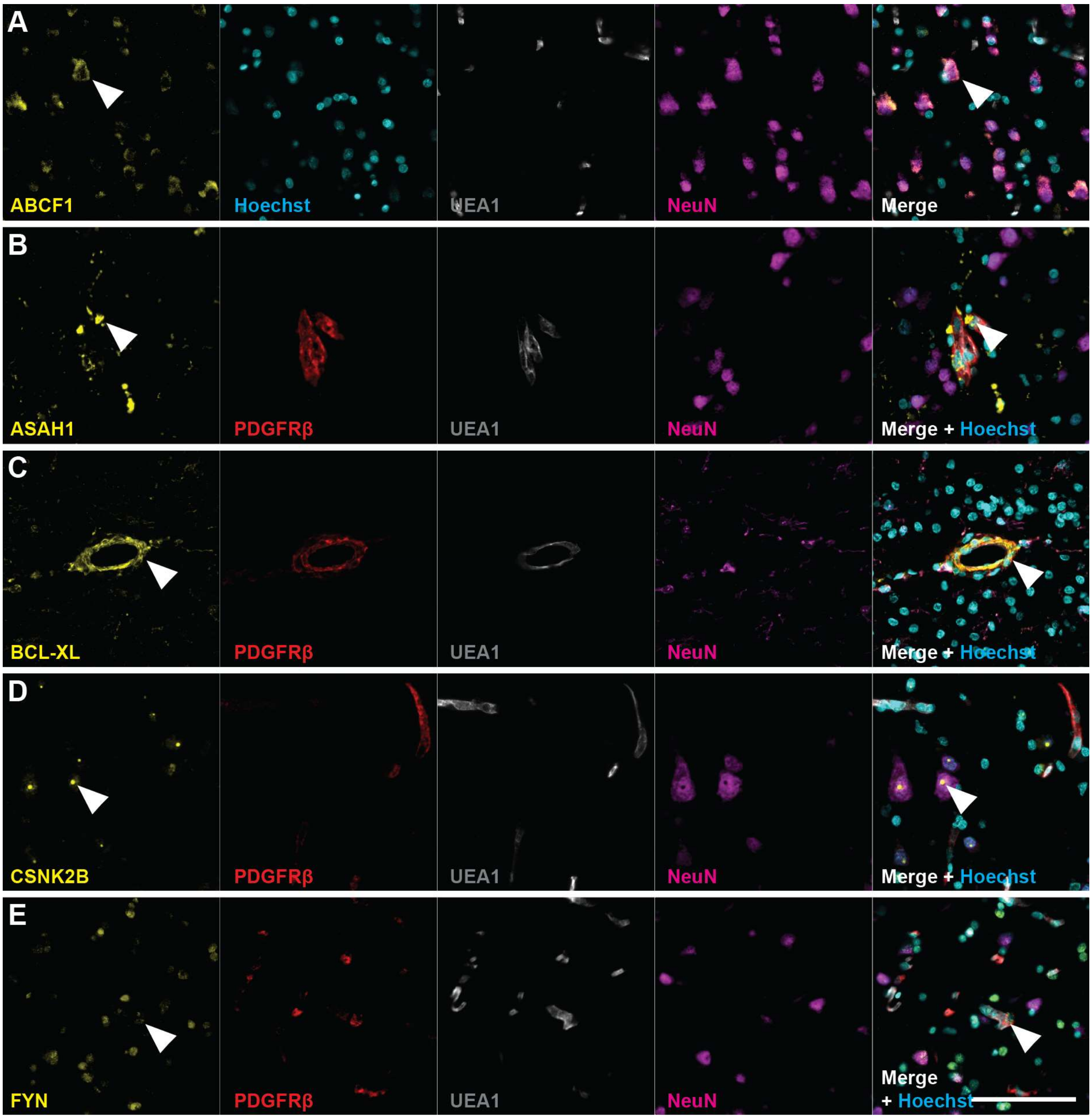


**Supplementary Figure 2. Single-channel images of DEG analogous proteins and cell-specific markers in PD MTG tissue.** White arrows indicate specific staining. Scale bar represents 100 μm.

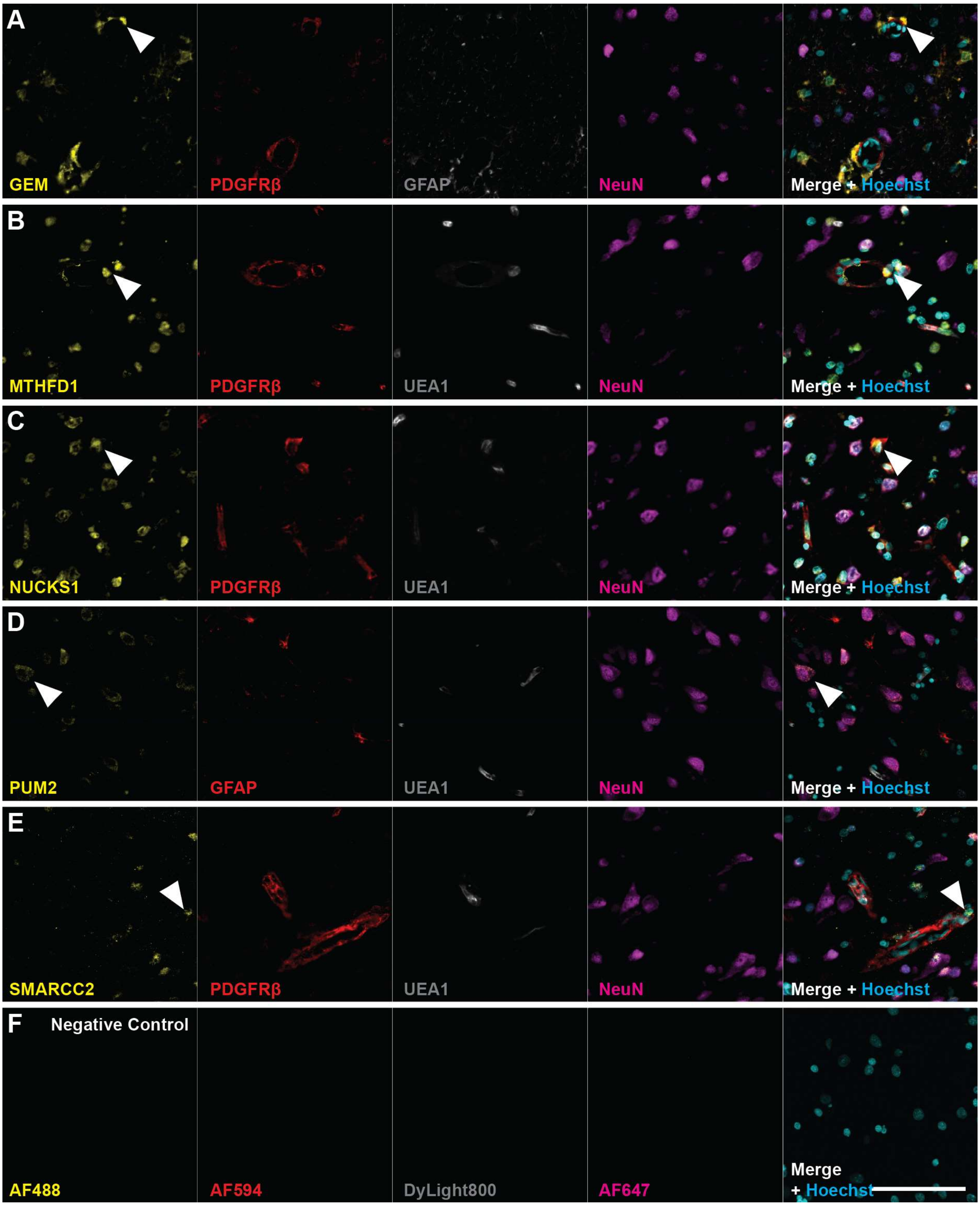


**Supplementary Figure 3. Single-channel images of DEG analogous proteins and cell-specific markers (A-E), and negative controls in PD MTG tissue (F).** White arrows indicate specific staining. Scale bar represents 100 µm.

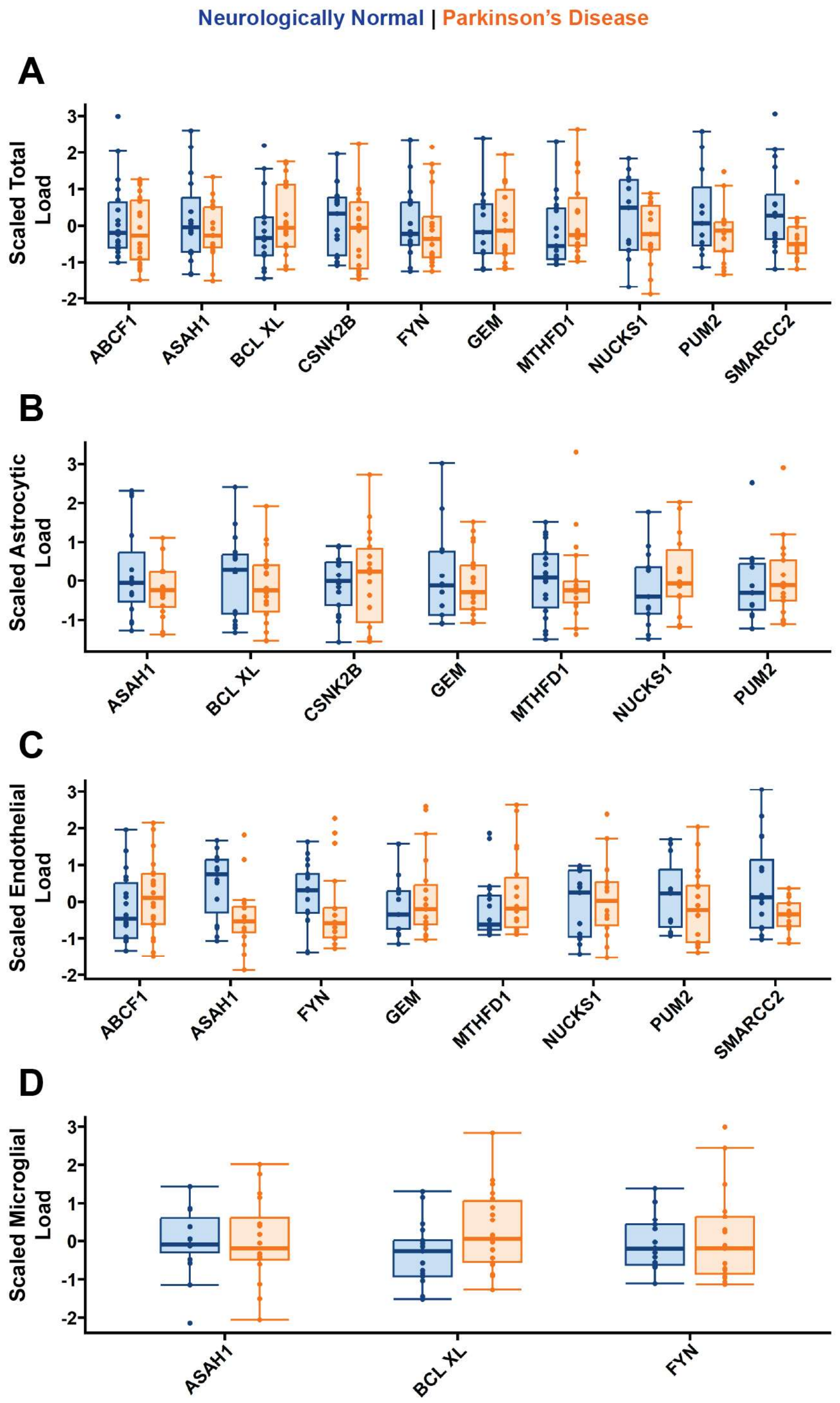


**Supplementary Figure 4. Total and Cell-specific expression of differentially expressed genes analogous proteins. (A-D)** No differences in total, nuclear, astrocytic and microglial protein expression were observed when comparing neurologically normal and PD cases. Normality of the datasets was evaluated using the Shapiro-Wilk test. Based on the results, either a t-test or a Mann-Whitney U test was performed to compare the neurologically normal and PD groups. Statistical significance was indicated as $p < 0.05$ (*) and $p < 0.01$ (**).

**Supplementary Table 7. Non scaled Mean total and cell-specific protein load for neurologically normal and Parkinson's disease cases.**

| | | Protein Load (%) | | | | | | |
|---|---|---|---|---|---|---|---|---|
| | | **Total** | **Pericytic** | **Nuclear** | **Neuronal** | **Microglial** | **Astrocytic** | **Endothelial** |
| **ABCF1** | *Neurologically normal (n =18)* | 3.1 ± 1.2 | - | 8.5 ± 3.2 | 17.5 ± 7.1 | - | - | 3.5 ± 1.3 |
| | *PD (n =20)* | 2.8 ± 1.0 | - | 8.1 ± 3.1 | 14.7 ± 5.7 | - | - | 3.9 ± 1.5 |
| **ASAH1** | *Neurologically normal (n = 15)* | 2.6 ± 1.1 | 12.9 ± 3.5 | 5.2 ± 1.8 | 12.0 ± 12.5 | 7.6 ± 2.0 | 2.7 ± 1.2 | 5.9 ± 1.7 |
| | *PD (n = 16)* | 2.3 ± 0.7 | 9.6 ± 3.8 | 4.6 ± 1.7 | 11.9 ± 11.0 | 7.7 ± 2.6 | 2.3 ± 0.7 | 4.3 ± 1.7 |
| **BCL-XL** | *Neurologically normal (n = 17)* | 2.9 ± 1.9 | **12.1 ± 9.2** | **4.4 ± 2.7** | - | 11.4 ± 6.2 | 7.3 ± 3.2 | - |
| | *PD (n =18)* | 3.5 ± 1.8 | **31.2 ± 14.6*** | **7.7 ± 3.4*** | - | 16.1 ± 8.2 | 6.4 ± 2.8 | - |
| **CSNK2B** | *Neurologically normal (n = 20)* | 1.0 ± 0.6 | - | 8.5 ± 4.5 | 10.1 ± 7.1 | - | 1.6 ± 0.8 | - |
| | *PD (n = 18)* | 0.8 ± 0.5 | - | 6.9 ± 4.8 | 6.3 ± 4.3 | - | 1.7 ± 1.0 | - |
| **FYN** | *Neurologically normal (n = 17)* | 2.3 ± 1.4 | 12.5 ± 5.7 | 21.0 ± 11.3 | 22.0 ± 22.2 | 7.7 ± 4.1 | - | 3.8 ± 1.7 |
| | *PD (n = 17)* | 2.2 ± 1.6 | 13.8 ± 8.2 | 21.4 ± 19.8 | 19.4 ± 18.8 | 8.8 ± 7.1 | - | 3.1 ± 2.2 |
| **GEM** | *Neurologically normal (n =13)* | 1.5 ± 1.3 | 10.3 ± 7.9 | 3.4 ± 2.4 | 1.3 ± 0.9 | - | 8.8 ± 9.1 | 2.8 ± 2.1 |
| | *PD (n = 19)* | 1.5 ± 1.2 | 10.7 ± 7.6 | 4.3 ± 3.0 | 1.1 ± 0.9 | - | 7.6 ± 6.0 | 3.6 ± 2.9 |
| **MTHFD1** | *Neurologically normal (n = 18)* | 0.6 ± 0.6 | 3.9 ± 2.5 | 5.7 ± 5.5 | - | - | 1.5 ± 0.9 | 1.0 ± 1.0 |
| | *PD (n =18)* | 0.8 ± 0.7 | 5.0 ± 3.0 | 7.6 ± 5.9 | - | - | 1.4 ± 1.0 | 1.5 ± 1.3 |
| **NUCKS1** | *Neurologically normal (n = 13)* | 11.7 ± 3.6 | 41.7 ± 23.7 | 52.4 ± 14.5 | **81.3 ± 7.8** | - | 22.4 ± 13.5 | 13.3 ± 6.1 |
| | *PD (n = 14)* | 10.2 ± 2.8 | 32.4 ± 16.1 | 55.2 ± 12.1 | **70.5 ± 12.3*** | - | 27.2 ± 14.1 | 14.5 ± 7.3 |
| **PUM2** | *Neurologically normal (n = 11)* | 2.1 ± 1.4 | - | 8.7 ± 6.9 | 17.1 ± 9.7 | - | 4.9 ± 3.6 | 22.1 ± 13.7 |
| | *PD (n = 17)* | 1.4 ± 0.8 | - | 6.7 ± 4.1 | 12.3 ± 7.3 | - | 5.2 ± 3.4 | 18.4 ± 13.6 |
| **SMARCC2** | *Neurologically normal (n = 16)* | 0.8 ± 0.4 | 2.1 ± 1.0 | 2.0 ± 1.0 | 1.9 ± 1.0 | - | - | 1.0 ± 0.6 |
| | *PD (n = 17)* | 0.5 ± 0.2 | 2.0 ± 0.6 | 1.3 ± 0.5 | 1.4 ± 0.6 | - | - | 0.7 ± 0.2 |

*Bolded values indicate significant differential expression between groups

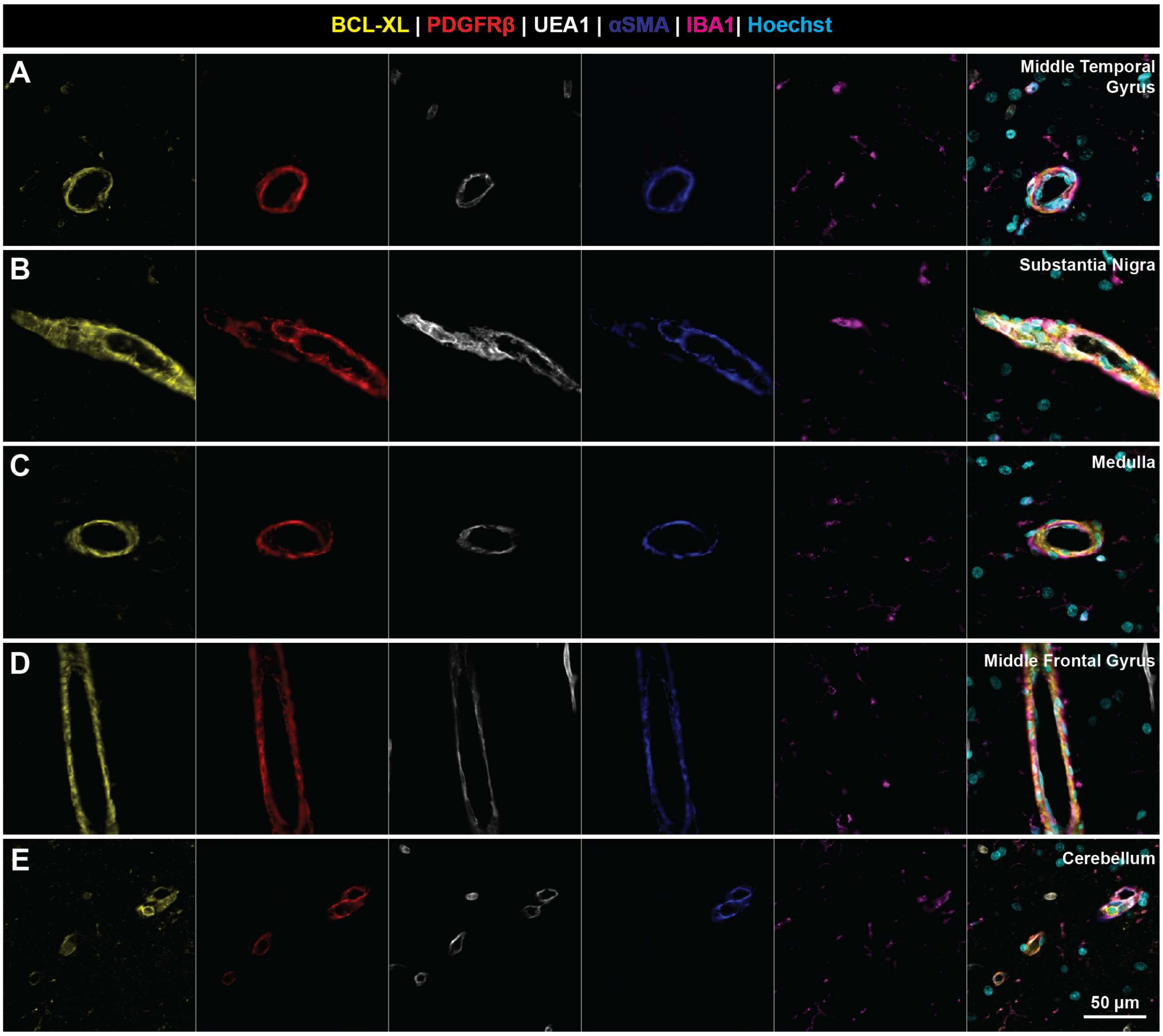


**Supplementary Figure 5. Single-channel images of BCL-XL and cell-specific markers across multiple PD brain regions.** The staining with BCL-XL, PDGFRß, UEA1, αSMA and IBA1, were overlayed to that with Hoechst in the panel most to the right for the A, Middle Temporal Gyrus, B, Substantia Nigra, C, Medulla, D, Middle Frontal Gyrus and E, Cerebellum. Staining's present from case PD65.

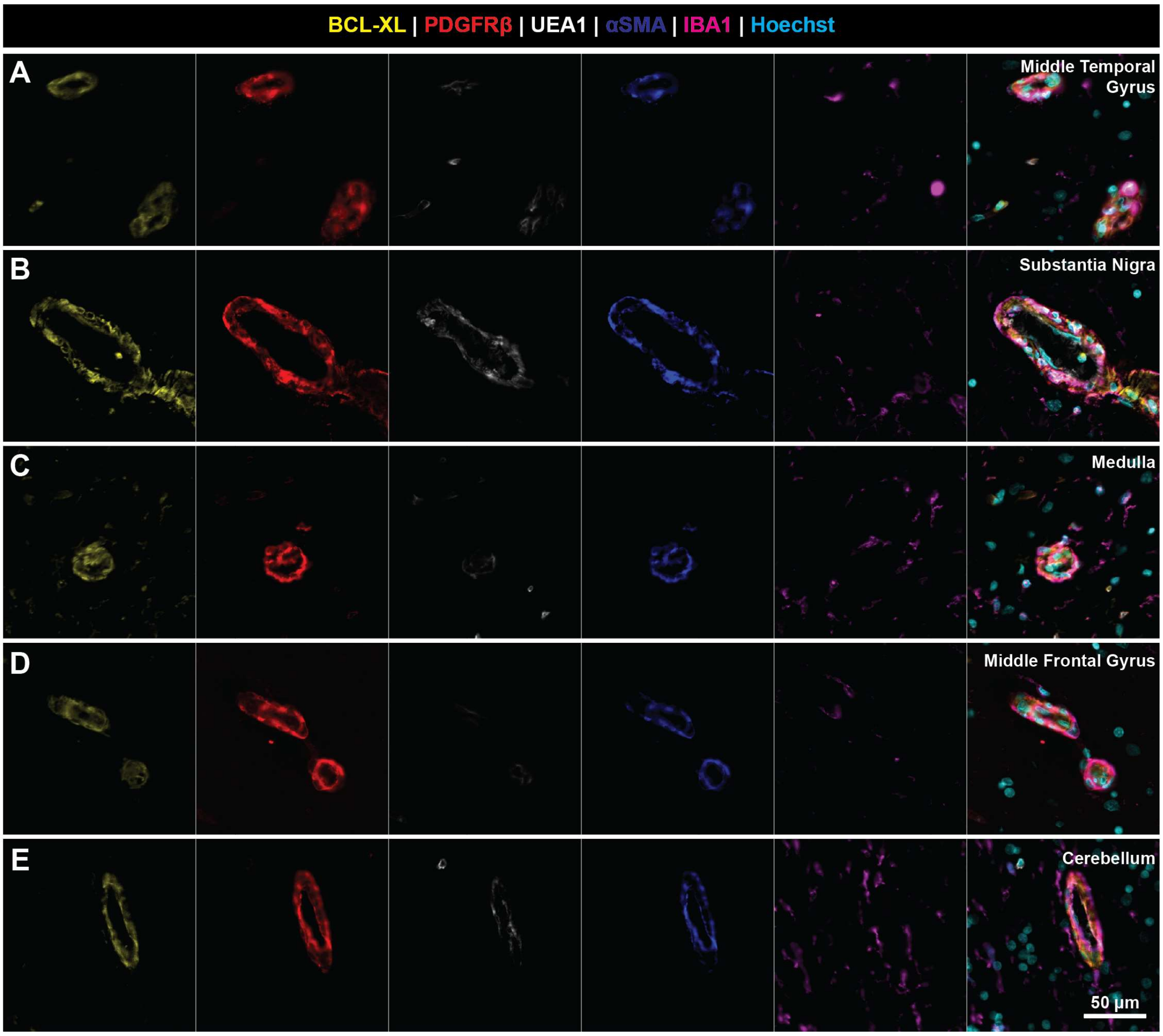


**Supplementary Figure 6. Single-channel images of BCL-XL and cell-specific markers across multiple neurologically normal brain regions.** The staining with BCL-XL, PDGFRß, UEA1, αSMA and IBA1, were overlayed to that with Hoechst in the panel most to the right for the A, Middle Temporal Gyrus, B, Substantia Nigra, C, Medulla, D, Middle Frontal Gyrus and E, Cerebellum. Staining's presented from case H239.

**Supplementary Table 8. Non-scaled BCL-XL expression values across multiple brain regions**

| | | BCL-XL Load (%) ± SD | | | | | |
|---|---|---|---|---|---|---|---|
| | | **Total** | **Pericytic** | **Nuclear** | **Microglial** | **Smooth Muscle** | **Endothelial** |
| **Middle Temporal Gyrus** | *Neurologically normal (n = 9)* | 0.3 ± 0.1 | **15.1 ± 3.1** | 0.9 ± 0.3 | 4.5 ± 2.5 | 20.4 ± 5.9 | 6.4 ± 5.5 |
| | *PD (n = 11)* | 0.5 ± 0.2 | **22.6 ± 6.8*** | 1.2 ± 0.4 | 6.9 ± 2.5 | 25.1 ± 6.1 | 9.3 ± 6.6 |
| **Substantia Nigra** | *Neurologically normal (n = 10)* | 0.7 ± 0.6 | **17.2 ± 7.2** | 2.2 ± 1.1 | **5.9 ± 2.9** | 21.0 ± 10.6 | 7.3 ± 4.1 |
| | *PD (n = 11)* | 0.9 ± 0.3 | **26.1 ± 4.2*** | 2.8 ± 1.1 | **10.5 ± 3.9*** | 28.7 ± 7.1 | 12.8 ± 7.8 |
| **Medulla** | *Neurologically normal (n = 7)* | 0.8 ± 0.2 | 27.9 ± 4.4 | 3.1 ± 0.7 | 13.4 ± 5.8 | 32.8 ± 5.4 | 10.1 ± 2.7 |
| | *PD (n = 9)* | 0.6 ± 0.3 | 26.9 ± 7.5 | 2.2 ± 0.8 | 6.8 ± 2.7 | 30.8 ± 6.8 | 9.9 ± 9.0 |
| **Middle Frontal Gyrus** | *Neurologically normal (n = 10)* | 0.5 ± 0.2 | 16.6 ± 5.3 | 1.4 ± 0.5 | 8.9 ± 4.9 | 22.2 ± 8.1 | 6.8 ± 3.8 |
| | *PD (n = 11)* | 0.4 ± 0.3 | 19.8 ± 11.2 | 1.2 ± 0.8 | 6.6 ± 3.6 | 25.4 ± 10.9 | 9.9 ± 11.6 |
| **Cerebellum** | *Neurologically normal (n = 10)* | 0.6 ± 0.3 | 18.4 ± 5.9 | 0.6 ± 0.4 | 6.1 ± 3.8 | 23.5 ± 6.9 | 8.2 ± 8.9 |
| | *PD (n = 11)* | 0.6 ± 0.2 | 19.1 ± 5.7 | 0.5 ± 0.2 | 6.3 ± 1.9 | 24.0 ± 8.6 | 7.4 ± 6.1 |

*Bolded values indicate significant differential expression between groups

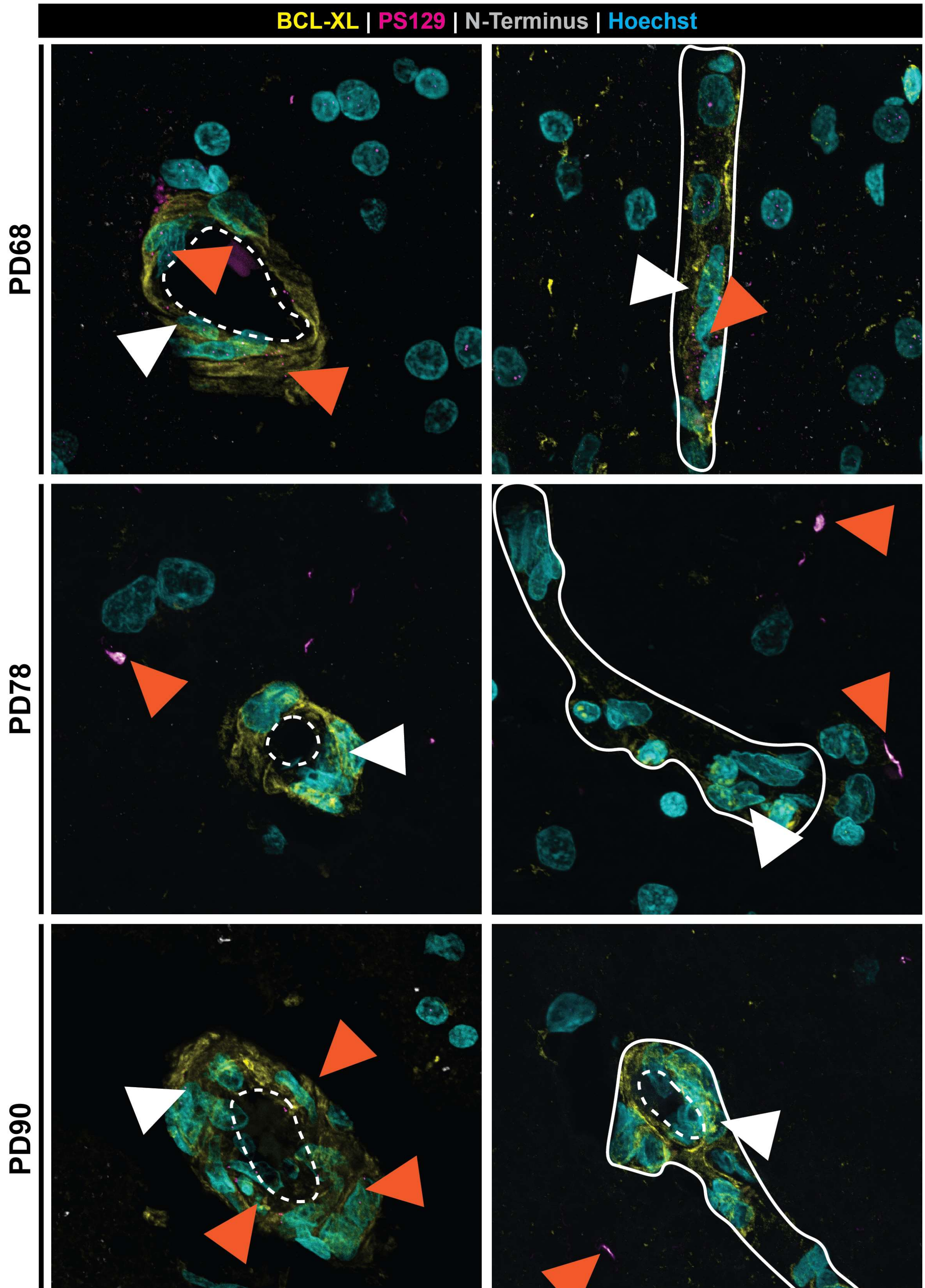
BCL-XL | PS129 | N-Terminus | Hoechst
PD68
PD78
PD90

**Supplementary Figure 7. Confocal Z-stack maximum projection immunofluorescence images illustrate no relationship between BCL-XL and α-Synuclein in the middle temporal gyrus of Parkinson's Disease cases.** White arrows indicate specific BCL-XL staining. Orange arrows indicate α-Synuclein staining. White dotted lines indicate vessel lumens. Solid white outlines delineate vessel borders Scale bar represents 30 μm.

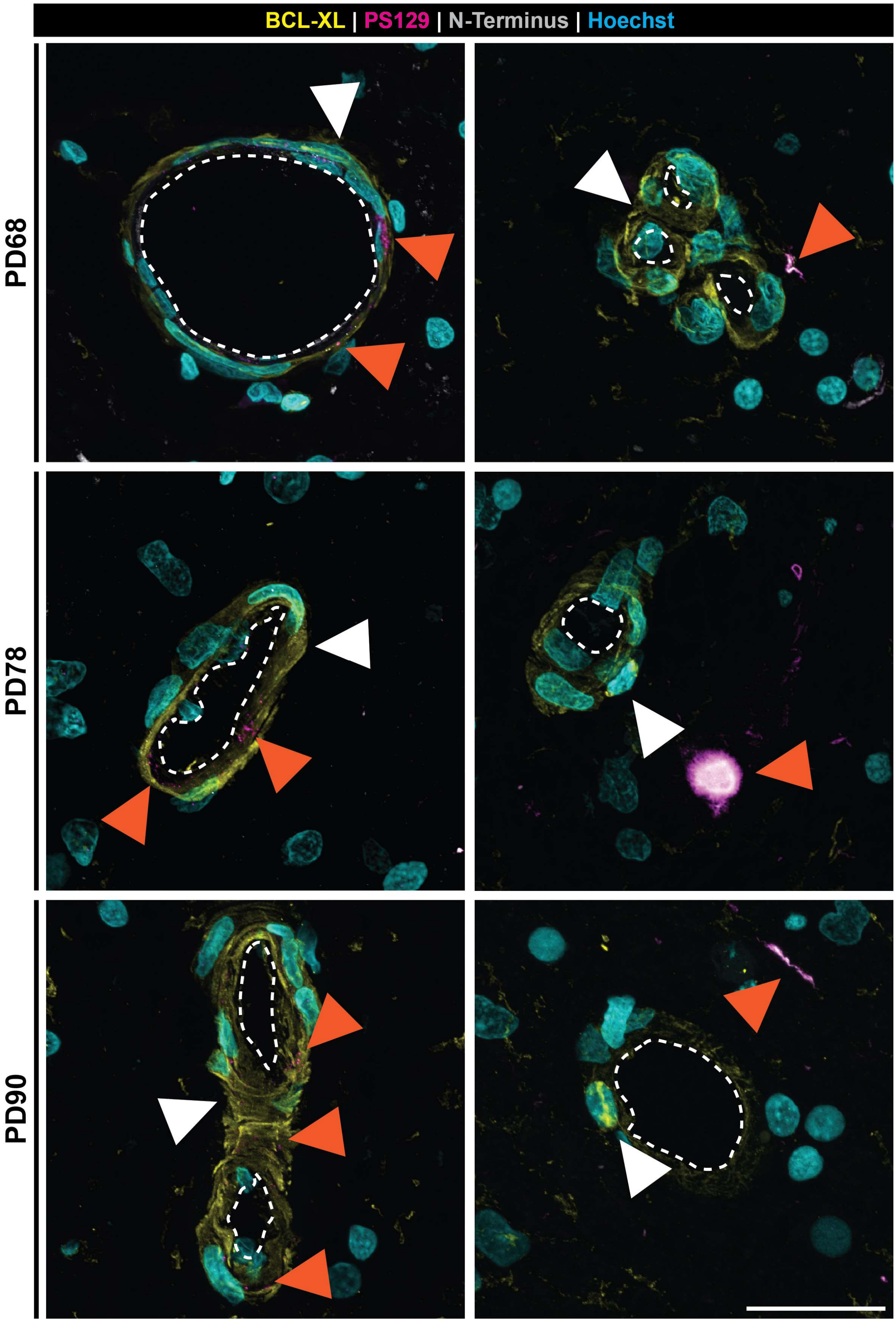

BCL-XL | PS129 | N-Terminus | Hoechst
PD68
PD78
PD90

**Supplementary Figure 8. Confocal Z-stack maximum projection immunofluorescence images illustrate no clear relationship between BCL-XL and α-Synuclein in the substantia nigra of Parkinson's Disease cases.** White arrows indicate specific BCL-XL staining. Orange arrows indicate α-Synuclein staining. White dotted lines indicate vessel lumens. Scale bar represents 30 μm.

**Supplementary Table 9. Correlations of BCL-XL expression and α-Synuclein load across multiple Parkinson's Disease brain regions**

| | Spearman Rho | | | | | |
|---|---|---|---|---|---|---|
| | **Total** | **Pericytic** | **Nuclear** | **Microglial** | **Smooth Muscle** | **Endothelial** |
| **Middle Temporal Gyrus** | 0.59 | 0.43 | 0.66 | -0.05 | 0.41 | 0.14 |
| **Substantia Nigra** | -0.5 | -0.35 | -0.24 | -0.18 | -0.27 | -0.69 |
| **Medulla** | 0.64 | 0.40 | 0.57 | 0.52 | 0.40 | 0.71 |
| **Middle Frontal Gyrus** | 0.59 | 0.43 | 0.66 | -0.05 | 0.41 | 0.14 |
| **Cerebellum** | 0.37 | -0.2 | 0.28 | -0.38 | -0.35 | -0.02 |

No correlations exhibited a p value < 0.05
